# Improving Recombinant Antibody Production Using FcBAR: An In Situ Approach to Detect and Amplify Protein-Protein Interactions

**DOI:** 10.1101/2025.06.12.659199

**Authors:** Mina Ying Min Wu, Frances Rocamora, Mojtaba Samoudi, Caressa M. Robinson, Chih-Chung Kuo, Nuša Pristovšek, Lise Marie Grav, Helene Faustrup Kildegaard, Gyun Min Lee, Alexandre Rosa Campos, Nathan E. Lewis

## Abstract

Recombinant proteins, in particular monoclonal antibodies and related molecules, have become dominant therapeutics. As they are produced in mammalian cells, they require the concerted function of hundreds of host cell proteins in the protein secretion pathway. However, the comprehensive set of host cell machinery involved remains unclear. Thus, it is often unknown why some recombinant proteins fail to express well. Here we present and deploy an approach called Fc-targeting Biotinylation by Antibody Recognition (FcBAR), which allows for the *in situ* detection of protein-protein interactions for any recombinant protein with Fc domain. Briefly, cells are permeabilized and incubated with an anti-Fc antibody, conjugated with horseradish peroxidase. All proteins interacting with Fc-bearing proteins are then biotinylated, pulled down and identified via mass spectrometry. We applied this method on a panel of rituximab-producing CHO-S clones with a range of productivity levels. Through analysis of FcBAR protein-protein interactions and RNA-Seq, we identified protein interactions positively correlated with rituximab secretion, and tested 7 of these targets. We found overexpression of AGPAT4, EPHX1, and NSDHL significantly increased rituximab production. Thus, FcBAR provides an unbiased approach to measure PPIs supporting recombinant antibody production *in situ*, and can guide efforts to boost production of biotherapeutics and biosimilars by addressing production bottlenecks.

## Introduction

The most high-demand biopharmaceutical products are monoclonal antibodies (mAbs), which accounted for more than half of total new pharmaceutical product approvals from 2015-2022 and dominate global sales, bringing in more than 60% of the sales in 2021 (Walsh & Walsh, 2022). The demand for mAbs and related biologics will continue to grow, to address their need for treating complex illnesses. Although biosimilars are making mAbs more affordable, the cost remains high. Moreover, some therapeutic proteins are difficult to express, sometimes due to bottlenecks in the secretory pathway, which limits their accessibility. Here, we use rituximab, a well-characterized mAb, as a model protein to investigate proteomic interactions that influence its secretion, with the goal of identifying strategies to enhance its production. Rituximab is a glycosylated murine/human chimeric IgG1κ mAb that targets the surface marker CD20 on malignant B-cells to treat non-Hodgkin’s lymphoma, chronic lymphocytic leukemia, and rheumatoid arthritis (Baldo, 2016; Selewski et al., 2010). Biosimilars of rituximab, such as Ruxience, have shown a 30% reduction in price compared to Rituxan (Kvien et al., 2022). Although biosimilars are making rituximab and other mAb treatments more affordable and accessible, the cost remains high.

One approach to reduce the cost of rituximab is to increase its production yield. Rituximab and its US-approved biosimilars are produced in Chinese Hamster Ovaries (CHO) cells. Optimizing the fed-batch bioprocess for rituximab-expressing CHO cells can increase in volumetric productivity (Mellahi et al., 2019). Building on these promising bioprocess results, our focus is on optimizing the secretory pathway machinery (SecM) at the cellular level to enhance rituximab productivity, as transgene mRNA and protein levels do not always correlate with high titers. This is because secretory pathway inefficiencies often act as bottlenecks in recombinant protein production.

The SecM involves over 1,157 genes associated with post-translational modifications (PTMs), protein folding, quality control (ER-associated degradation pathway, Unfolded Protein Response), and trafficking between the endoplasmic reticulum (ER) and Golgi apparatus in the mammalian system (Masson et al., 2024b). Given that many of the SecM proteins will physically interact with a secreted protein of interest to facilitate its folding, modification and transport through the pathway, co-expressing certain SecM components with difficult-to-express proteins in CHO cells has improved secretion (Malm et al., 2022). For mAbs, the bottleneck is primarily at the folding and assembly stages (Dinnis & James, 2005; Kaneyoshi et al., 2019). Overexpression of vesicle-associated proteins such as SNAP-23 and VAMP, which facilitate vesicle docking and fusion, has led to increased production of recombinant proteins including anti-CD18 monoclonal antibodies (IgG1) (Peng et al., 2011). However, the host cell machinery required for specific secreted proteins remains unclear, so it is difficult to know what should be augmented for any given cell line or product.

To address this issue, here we present FcBAR, an unbiased approach adapted from the Biotinylation by Antibody Recognition (BAR) method (Bar et al., 2018). Fc-BAR allows one to quantify protein-protein interactions (PPIs) between Fc-bearing proteins (e.g., mAbs, Fc-fusion proteins, etc.) and the host-cell proteome. Briefly, in FcBAR, the cell line of interest is fixed, permeabilized, and incubated with an anti-Fc antibody conjugated with horseradish peroxidase (HRP). All interacting proteins are then biotinylated by treatment with H_2_O_2_ and tyramide-biotin; biotinylated proteins are then pulled down and analyzed by mass spectrometry (MS). Unlike other methods, such as BioID, which requires the addition of a biotin ligase domain to the protein of interest (POI), our FcBAR approach can be done on any cell line without modifying the host or POI (Kim et al., 2016; Pfeiffer et al., 2022; Sears et al., 2019). We hypothesize that FcBAR can help identify known and novel protein interactors in the secretory pathway, which represent SecM proteins that support mAb production. This approach is particularly relevant given that approximately 40% of heavy chains and 65% of light chains are retained in the ER and Golgi apparatus (Kaneyoshi et al., 2019).

We applied FcBAR to 10 rituximab-expressing clones with a range of productivity levels. We detected >1400 unique proteins that correspond to rituximab-expressing clones and are significant in the control. Furthermore, through a multiomics analysis of the FcBAR proteomics and RNA-Seq data, we compiled a list of protein interactions correlated with high productivity. Our results demonstrate that FcBAR is an effective tool for identifying proteins in the secretory pathway that enhances production yield. This multi-omics approach shows promise for addressing production challenges for rituximab and potentially other difficult-to-express proteins.

## Results

### Generating Stable Cell Lines Expressing Rituximab

A plasmid containing genes for the heavy chain (HC) and light chain (LC) of rituximab was randomly integrated into suspension CHO-S cells, and the cells were subsequently cloned by single cell sorting. The protein levels of rituximab were quantified for each clone through the Octet RED96 system, a bio-layer interferometry instrument that uses surface plasmon resonance (SPR) technology to measure protein interactions. The clones were sorted into low or high producers based on the amount of rituximab they produced (pg per cell per day). We identified 10 clones with similar mRNA levels but varying rituximab production levels (Supplementary Figure S1B). Three low producers—B2, E9, and G12—had productivity ranging from 6.50 to 7.20 pg/cell/day, while seven high producers—A4, A6, B5, E6, E7, E10, and F1— showed productivity ranging from 8.87 to 14.46 pg/cell/day (Figure 1A). Expression levels of rituximab were quantified by SPR due to its sensitivity and suitability for measuring both low-abundance and highly abundant proteins. Additionally, Western blot analysis confirmed successful secretion of rituximab in each clone (Supplementary Figure S1A).

**Figure 1.**
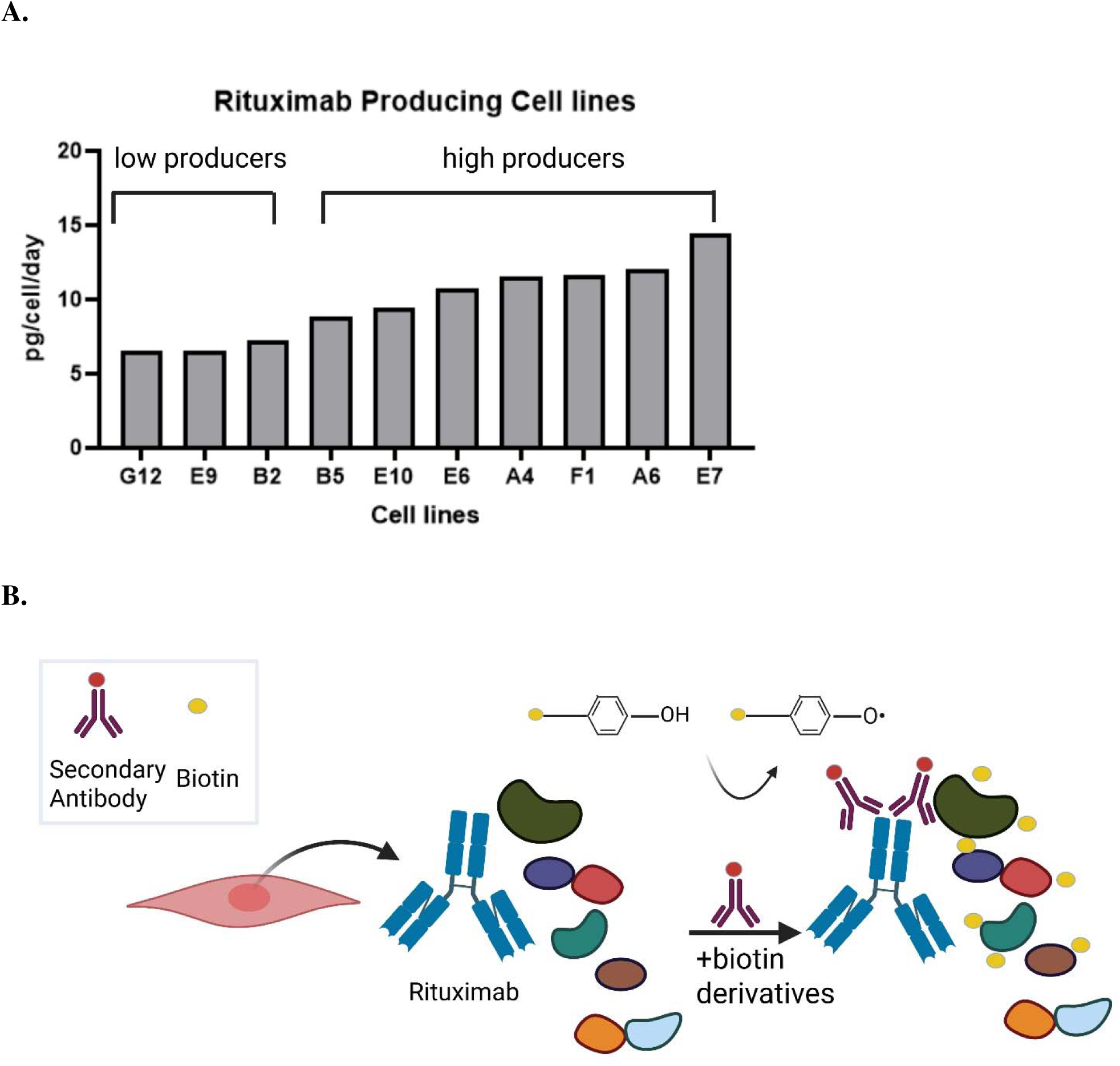
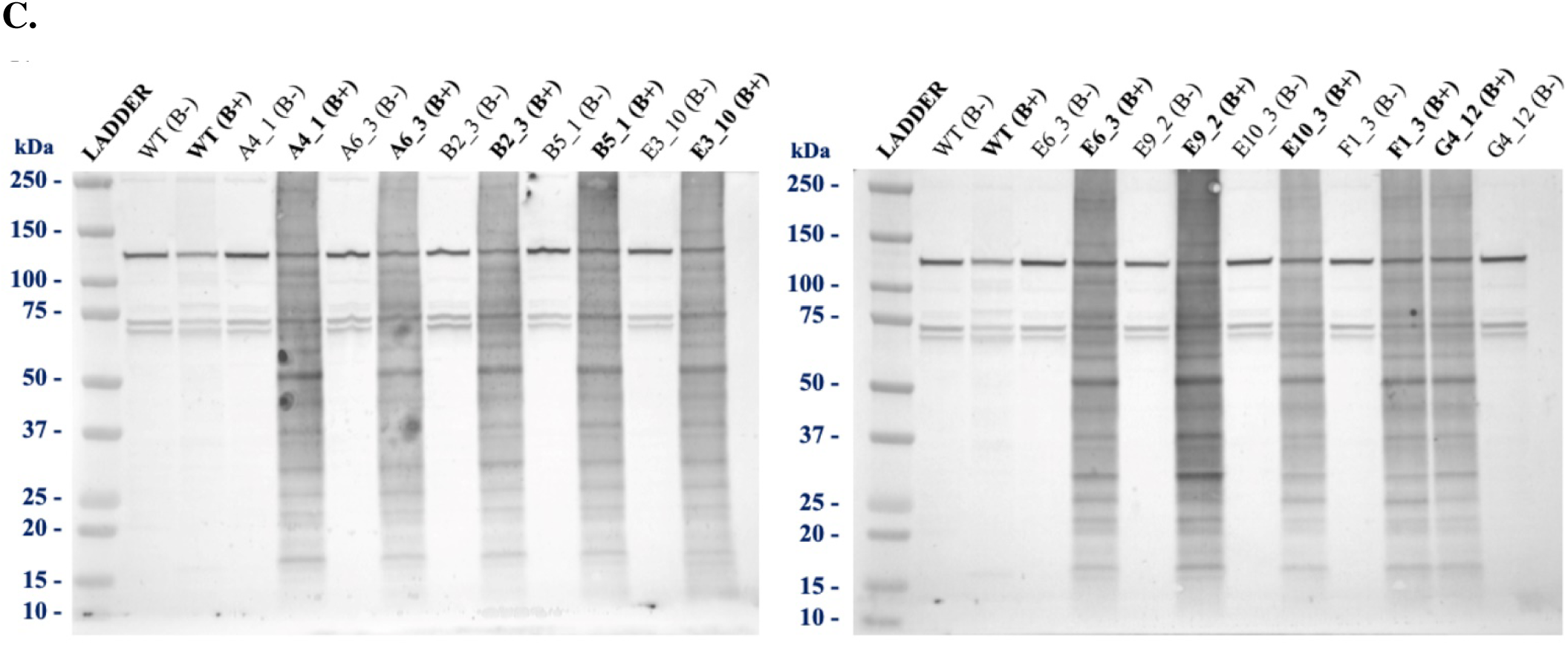
Utilizing biotinylation by antibody recognition targeting the Fc Region (FcBAR) to study protein-protein interactions in rituximab clones. **(A)** Surface Plasmon Resonance (SPR) identified three low-producing rituximab clones (G12, E9, and B2) with productivity ranging from 6.50 to 7.20 pg/cell/day, and seven high-producing rituximab clones (B5, E10, E6, A4, F1, A6, and E7) with productivity ranging from 8.87 to 14.46 pg/cell/day. **(B)** An illustrated depiction of the FcBAR method applied to rituximab-expressing clones. Following fixation and permeabilization, cells were stained with an HRP-conjugated anti-human Fc antibody. Biotinylation was performed for 5 minutes, labeling proteins in proximity to rituximab. **(C)** Biotinylation was confirmed via Western blot using streptavidin-HRP staining. Each clone included a negative control (B-) in which biotin was not added. A CHO-S wild-type not expressing rituximab served as an additional control. Clones expressing rituximab and treated with biotin contained more biotinylated proteins, indicating successful biotinylation of proximate proteins associated with rituximab.

### Targeting the Fc Region of Rituximab using FcBAR

To investigate the PPIs of rituximab, FcBAR was optimized to target the Fc region of rituximab. FcBAR and bulk RNA-Seq were performed on the rituximab-expressing clones, harvesting the cells during the mid-exponential phase. FcBAR, the cells were fixed, permeabilized, and stained with an anti-human Fc antibody conjugated to HRP (Figure 1B). The cells were then biotinylated for 5 minutes, after which cell lysates were harvested. The biotinylated proteins were captured using streptavidin beads, eluted, and trypsinized for liquid chromatography-tandem mass spectrometry (LC-MS/MS) analysis.

Western blot analysis confirmed biotinylation of the samples, revealing the biotin profile by staining with streptavidin (Figure 1C). Each clone included a negative control (B-), where biotin was not added. The Western blot results showed that the rituximab-producing clones contained more biotinylated proteins compared to both the WT control and the negative control, confirming the specificity of the endogenous biotin labeling for interacting proteins. Additionally, the cells were fixed, permeabilized, and stained with an anti-human Fc-650 antibody targeting rituximab, DAPI for nuclear staining, and streptavidin-594 for biotinylated proteins. Confocal microscopy verified the co-localization of rituximab detected in each rituximab producing clone in the far-red channel (650 nm; shown in red) with biotinylated proximal proteins detected in the red channel (594 nm; shown in green) (Figure 2A). This was quantitatively confirmed with Pearson’s coefficient of 0.87, indicating strong colocalization between the rituximab and the biotinylated proximate proteins (Figure 2B).

**Figure 2:**
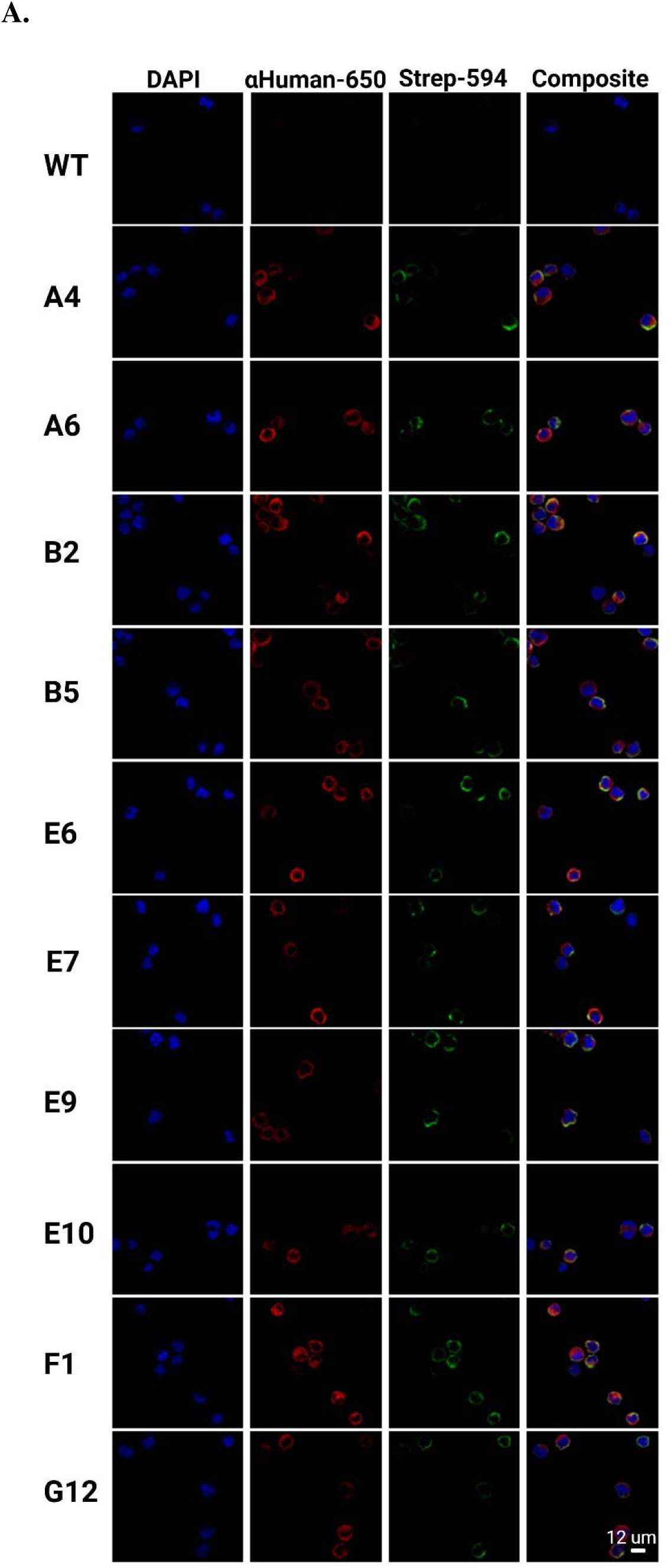
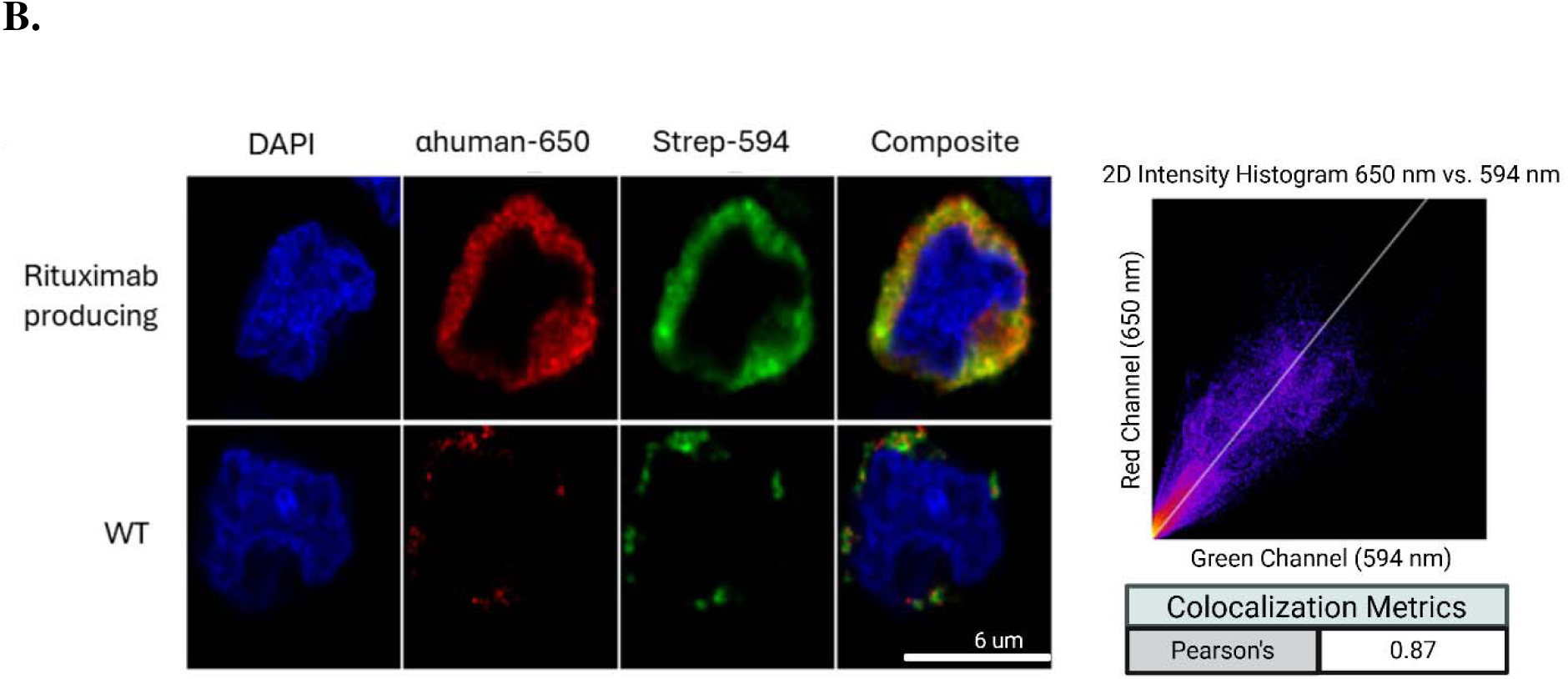
Colocalization of rituximab with biotinylated proteins confirmed by confocal microscopy after BAR. **(A)** Confocal microscopy of the BAR samples for each rituximab clone demonstrated co-localization of rituximab (far red channel, anti-human-650 nm shown in red) with biotinylated proximate proteins (red channel, streptavidin-594 nm shown in green). The scale bar corresponds to 12 um. **(B)** Intensity plot between the 650 (rituximab) and 594 (streptavidin) channels showed co-localization with a Pearson’s coefficient of 0.87, indicating strong colocalization between the rituximab and the biotinylated proximate proteins. Cells were fixed, permeabilized, and stained with DAPI for nuclear staining, anti-human Fc-650 nm antibody targeting rituximab, and streptavidin-594 nm to detect biotinylated proteins. The scale bar corresponds to 6 um.

### Preprocessing and Quality Control Analysis of FcBAR Proteomic Data

The FcBAR proteomic mass spectra were analyzed using MaxQuant software (Tyanova et al., 2016), and the MS/MS spectra were searched against the *Cricetulus griseus* Uniprot protein sequence database. To analyze the proteomic data, we utilized the DEP package (Zhang et al., 2018) in RStudio, first performing quality control (QC) by plotting the total protein abundance for each sample. We expected the total amount of biotinylated proteins to be similar across replicates for each sample. Based on the plot of biotinylated proteins detected per sample, one replicate each from cell lines B5 and E7 showed significantly lower numbers of biotinylated proteins and thus failed quality control; these replicates were excluded from further analysis (Figure 3A).

**Figure 3.**
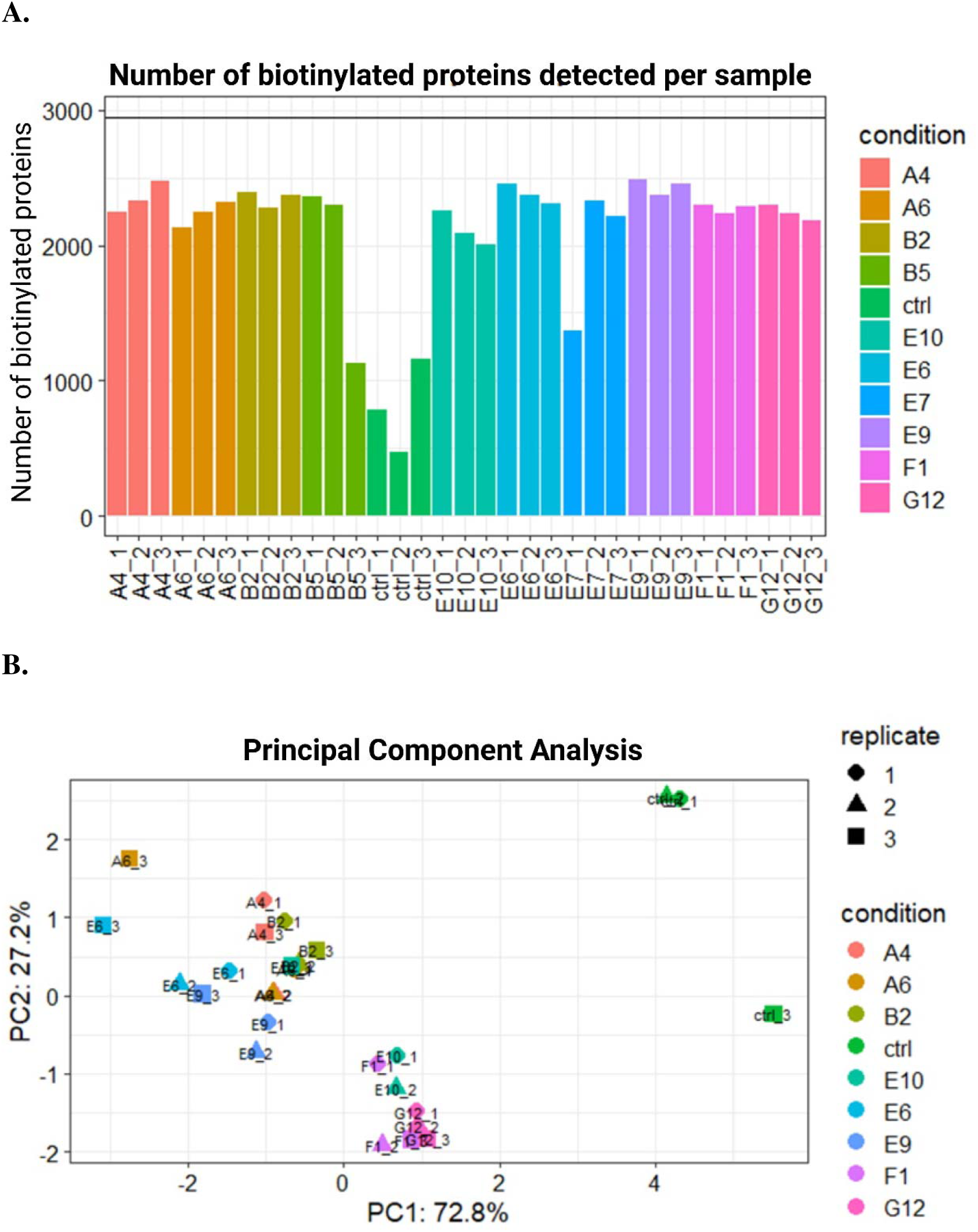
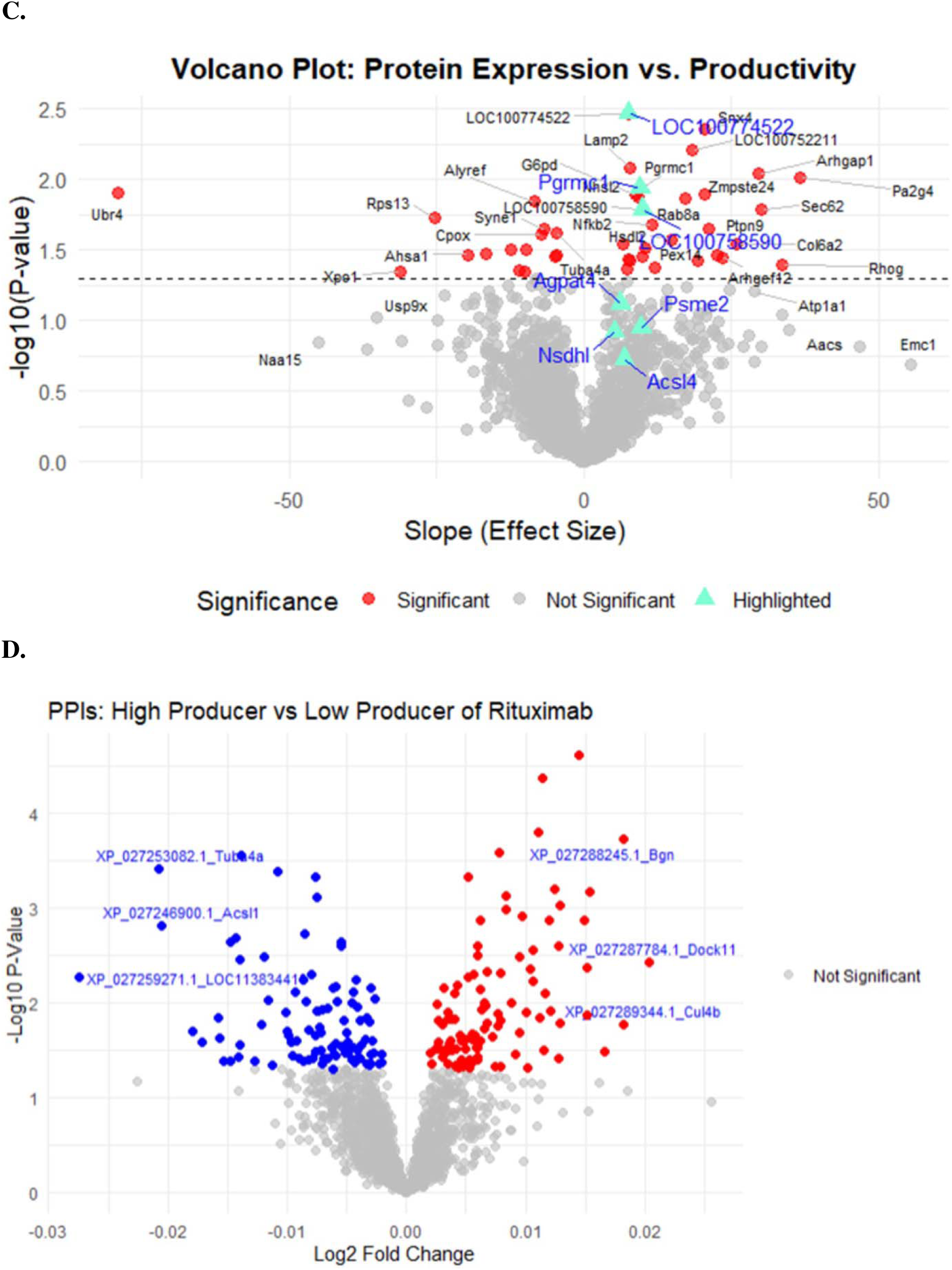
Quality control of FcBAR and comparative proteomics analysis: linear regression of protein abundance versus productivity and differential expression analysis between high- and low-productivity clones. **(A)** The rituximab FcBAR samples were plotted against the number of biotinylated proteins detected per sample. During LC-MS/MS analysis, one replicate from each of clones B5 and E7 showed inconsistent numbers of captured biotinylated proteins compared to their respective replicates and was therefore excluded from the analysis. **(B)** PCA on the normalized proteomic dataset revealed distinct clustering, with non-producer cells (control) forming a separate cluster from the rituximab producer cell lines, as captured in PCA1, explaining 72% of the variance. **(C)** Linear regression analysis was performed on the protein abundance vs productivity (pg/cell/day) to identify proteins that show a positive association with high productivity (slope > 0, p < 0.05, and R² > 0.50). **(D)** Welch’s t-test was performed to compare high versus low producer cell lines, identifying 99 (red dots) and 101 (blue dots) proximate proteins as differentially expressed in the high and low producing cell lines, respectively (p < 0.05).

We normalized the data using a variance stabilizing transformation (VST), which transforms the data so that the variance of protein expression values is stabilized across the measurement range. In raw proteomic data, low-abundance proteins tend to have small variations (low variance), while high-abundance proteins tend to have larger variations (high variance), meaning that the variance in the data depends on the abundance. This makes it difficult to perform statistical comparisons (e.g., t-tests, ANOVA), and clustering methods may be biased. However, applying VST stabilizes the variance across all expression levels and allows for fair statistical comparisons. Next, the proteomic data was imputed using a left-shifted Gaussian distribution based on the assumption that missingness is not at random (MNAR), which means that missing values are likely to represent low-abundance proteins. This approach avoids false zeros by acknowledging that missing proteins are likely just below the limit of detection. It ensures that imputed values are realistic and improves the reliability of statistical tests for differential expressions. We then analyzed the variance and clustering of the rituximab clones using Principal Component Analysis (PCA), which transforms high-dimensional data into a lower-dimensional subspace while preserving the variance. The PCA plot clearly revealed that the nonproducers clustered separately from the producer cell lines, with PC1 capturing 72% of the total variance, indicating a clear difference between proteins captured in producer and nonproducer clones (Figure 3B).

### Identifying Protein Interactors with Proteomic and Transcriptomic Analysis

To identify proteins associated with productivity levels, we performed linear regression analysis using the mean expression values of each protein against the productivity (pg/cell/day) of each clone (Figure 3C). We identified 28 proteins with a significant positive association with productivity (p < 0.05, slope > 0, R^2^ > 0.50) (Supplementary Tables S1A and S3), which are related to protein trafficking/secretion (*Sec62, Atp11c, Tab8a, Snx4*), metabolic support (*LOC100774522* and *LOC100752211* relate to cytochrome b5 functions) and autophagy/lysosomal clearance (*Lamp2, Zmpste24*), suggesting these proteins may enhance rituximab secretion. In contrast, 15 proteins showed a significant negative association with productivity (p < 0.05, slope < 0, R² > 0.50) (Supplementary Tables S1B and S3), indicating potential inhibitory effects on productivity, which involve protein degradation (*Ubr4, Psmd11, Xpo11*) that may degrade essential proteins, ribosomal machinery (*Rps13, Alyref*) that may cause reduced translation capacity, and pro-apoptotic signals (*Stk24, Crlf1*) that may trigger cell cycle arrest or death, potentially indicating inhibitory factors.

In addition, we identified differentially expressed proteins by employing Welch’s t-test on the FcBAR data comparing high and low producer cell lines. We identified >1,000 potential interactors with rituximab, revealing 99 interactors significantly expressed (p < 0.05) and unique in high producers and 101 interactors significantly expressed (p < 0.05) and unique in low producers (Figure 3D). To complement the proteomic analysis, bulk RNA-Seq analysis was performed to determine if any of the identified PPIs overlapped with the transcriptomic data. Using Welch’s t-test with the DESeq2 package in RStudio, approximately 1300 genes were significantly expressed in high producers (p < 0.05) and >900 genes were significantly expressed in low producers (p < 0.05) at the mRNA level (Figure 4A).

**Figure 4.**
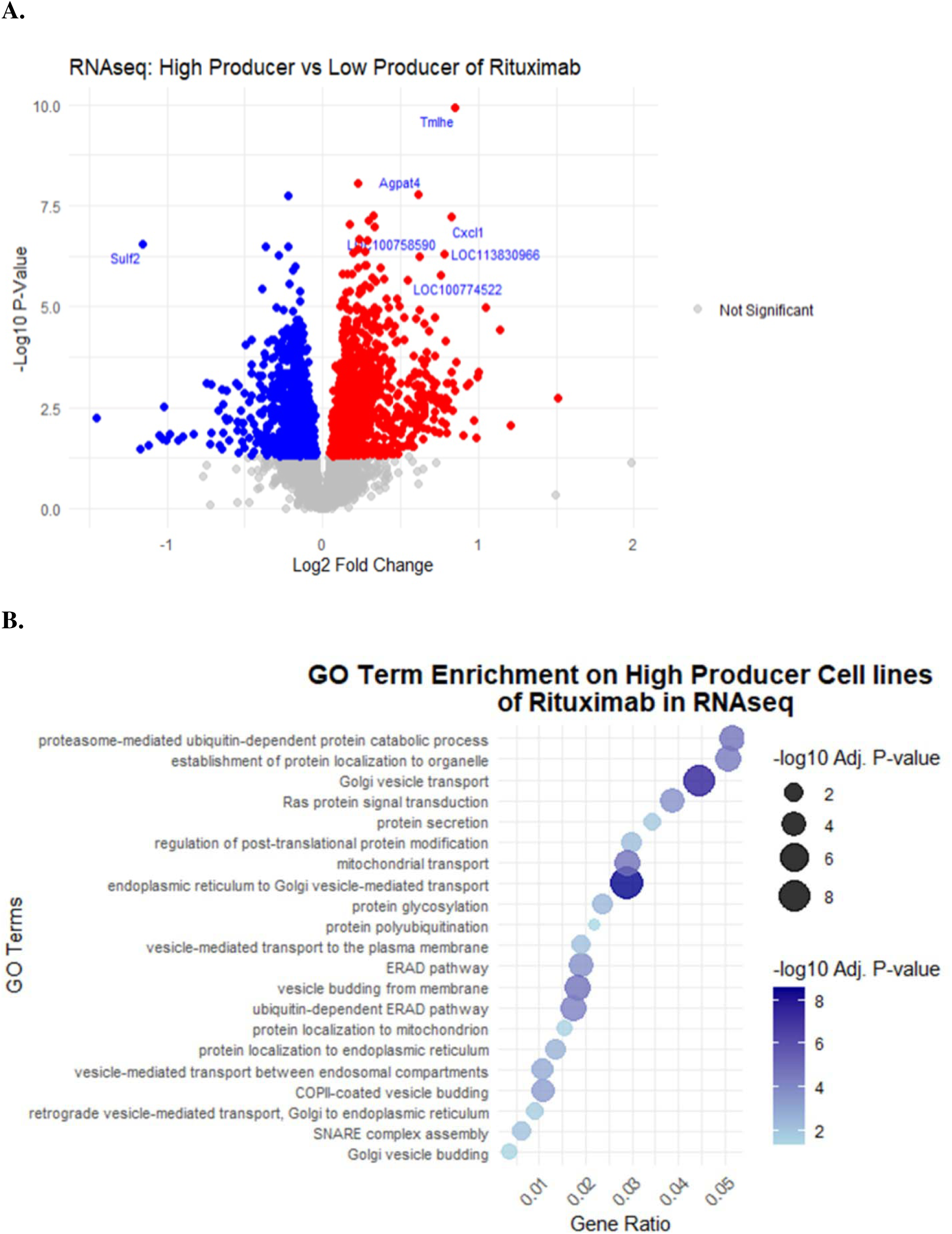
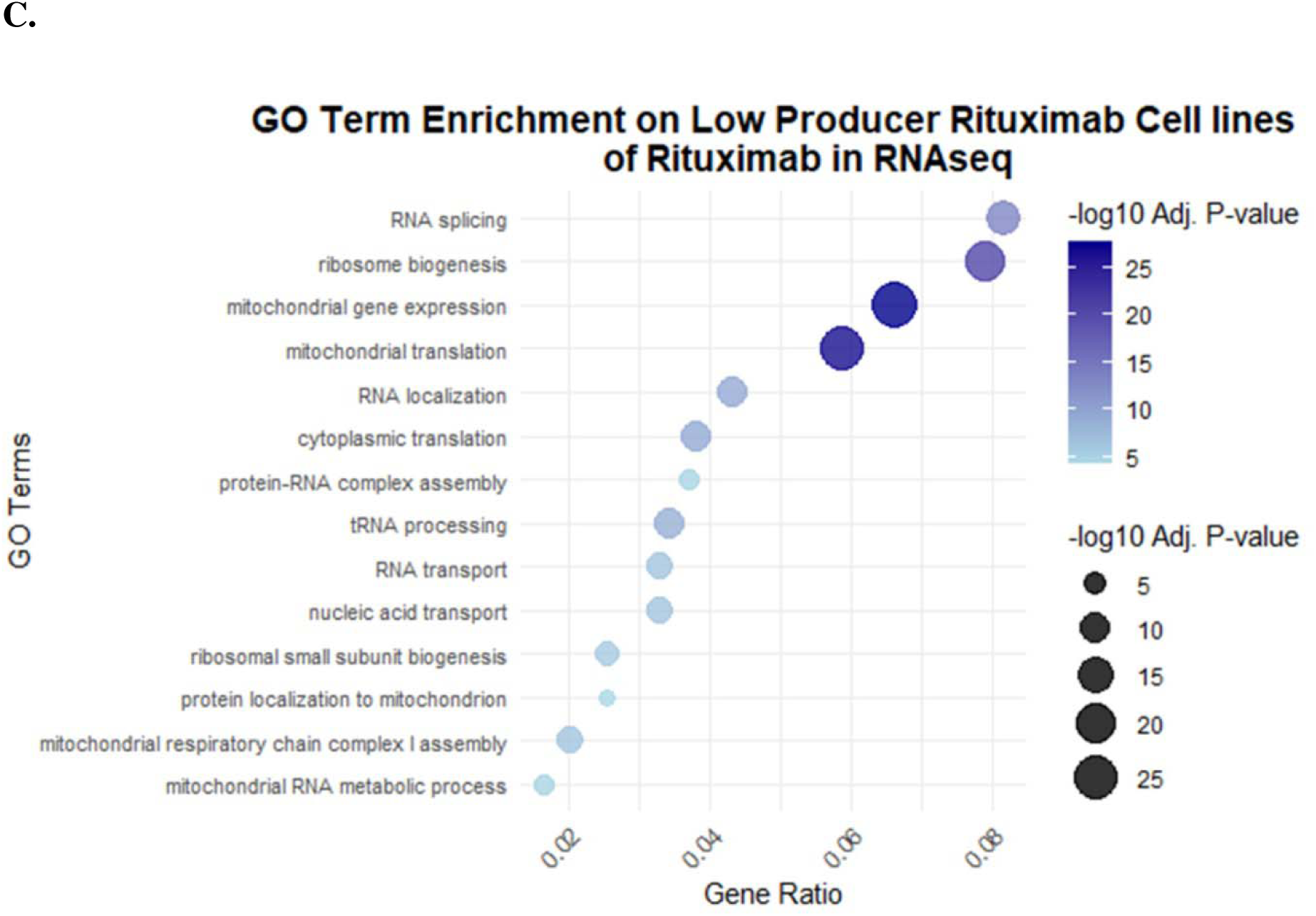
Bulk RNA-Seq transcriptomics in high- and low-producing rituximab cell lines. **(A)** Bulk RNA-Seq analysis was conducted to compare high- and low-producing cell lines using the DESeq2 package in RStudio. The analysis identified approximately >1,300 differentially expressed genes in high-producing cell lines (red dots) and >900 differentially expressed genes in low-producing cell lines (blue dots) (Welch’s t-test, p < 0.05). **(B)** Gene enrichment analysis focused on biological processes revealed significant pathways associated with the secretory pathway in high-producing cell lines. These pathways included protein secretion, ERAD pathway, vesicle budding, and related processes. **(C)** In contrast, low-producing cell lines showed gene enrichment primarily in RNA processing, splicing, modifications, and mitochondrial functions.

Gene enrichment analysis using the enrichGO function in the clusterProfiler package in R revealed that high-producing cells were enriched in biological processes related to the secretory pathway, such as ER-to-Golgi transport, ER stress, protein folding, and COPII vesicle budding (Figure 4B). In contrast, the significant genes in low producers were mainly related to RNA splicing/processing/modification/localization, mitochondria functions, translation, and protein synthesis.

We identified 68 proteins that exhibited significantly different abundances in PPIs and significantly different mRNA expression levels when comparing high-producing to low-producing clones (p < 0.05) (Supplementary Tables S2, S4, and S5). These proteins fell into several functional categories of interest. For example, some significant targets of particular interest included those with structural and cytoskeletal functions (e.g., *Pls3, Rdx, Sptan1*), Vesicle transport: *Dock11, Rab8a, Ocr1, Rtn4, Agpat4*), involvement in protein folding and trafficking (*Bcap31, Cul4b, Sacm1l, Sec62, Ergic1, Pgrmc1, UBE2N*), cell adhesion and extracellular matrix components (*Bgn, Col6a1/4a1/4a2, Thbs1/s3, Emilin1*), metabolism and energy production (*G6pd, Idh3g, Prps1, Dhcr7, Nsdhl*), and DNA/RNA processing (*Fmr1, Morc4, Sun2, Yars, Aars*).

From these significant genes, we selected seven of the most significant to test to see if their overexpression could increase rituximab production: AGPAT4, ACSL4, NSDHL, PSME2, LOC100774522 (UBE2N), PGRMC1, and LOC100758590 (EPHX1). These were of particular interest since they are anticipated to serve distinct but complementary cellular functions. AGPAT4 regulates membrane curvature (Karagiota et al., 2022), while ACSL4 catalyzes the activation of fatty acids to form acyl-CoA, a crucial step in lipid biosynthesis and fatty acid degradation. ACSL4 shows particular specificity for polyunsaturated fatty acids such as arachidonic acid and is enriched in ER subregions that form physical contacts with mitochondria, where it overlaps closely with the ER chaperone calnexin (Radif et al., 2018; Sen et al., 2020). Similarly, NSDHL participates in lipid droplet formation and cholesterol biosynthesis (Cunningham et al., 2009). In contrast, PSME2 (PA28β) functions in protein quality control as a subunit of the PA28 proteasome activator complex, which interacts with the 20S proteasome to degrade misfolded proteins (Rechsteiner & Hill, 2005). Together with PA28α, PSME2 forms a hexameric structure that facilitates HSP90-dependent refolding of denatured proteins such as luciferase (Minami et al., 2000). UBE2N also contributes to protein quality control as a member of the ubiquitin-conjugating enzyme family, working in conjunction with E1 (ubiquitin-activating enzyme) and E3 (ubiquitin ligase) enzymes to tag proteins for proteasomal degradation (Plafker et al., 2018; Valimberti et al., 2015). Additionally, PGRMC1 is an ER transmembrane protein that participates in ER-phagy by acting as a size-selective cargo receptor, targeting smaller, misfolded prohormones for degradation through its interaction with reticulon-3 (Chen et al., 2021). Finally, EPHX1 is an ER-localized enzyme involved in hydrolyzing endogenous epoxy fatty acids (Gautheron & Jéru, 2021; Václavíková et al., 2015). Collectively, these proteins were selected based on their essential roles in lipid metabolism, protein modification, and intracellular trafficking—processes critical for efficient protein secretion.

The three proteins, UBE2N, PGRMC1, and EPHX1 not only showed increased mRNA abundance and PPIs in the high producers, but they further showed significant positive associations with productivity (p < 0.05, R^2^ > 0.60). Thus, we tested the ability for overexpression of these 7 proteins to increase rituximab production.

### Overexpression of Identified Protein Interactors Increased Rituximab Expression

We tested selected targets by overexpressing the genes using PEI Max transient transfection in the low producer rituximab cell line, E9. Each test was conducted in triplicate. RNA was extracted on day 2 post-transfection for quantitative PCR (qPCR) confirmation of overexpression. All genes were successfully overexpressed, with Ephx1 showing particularly high expression, indicating that this protein is natively expressed at low levels in the cell line (Figure 5A).

**Figure 5.**
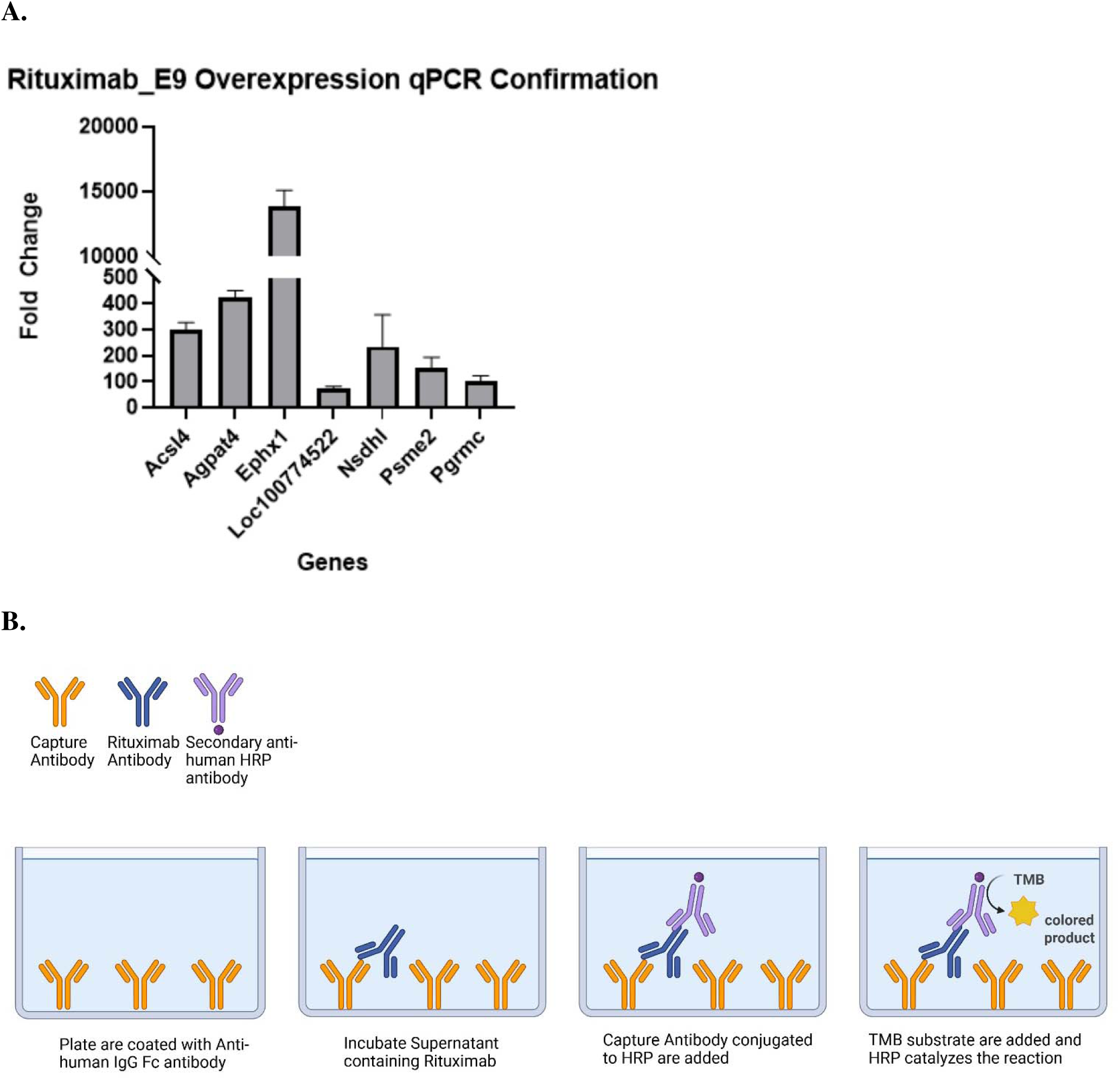
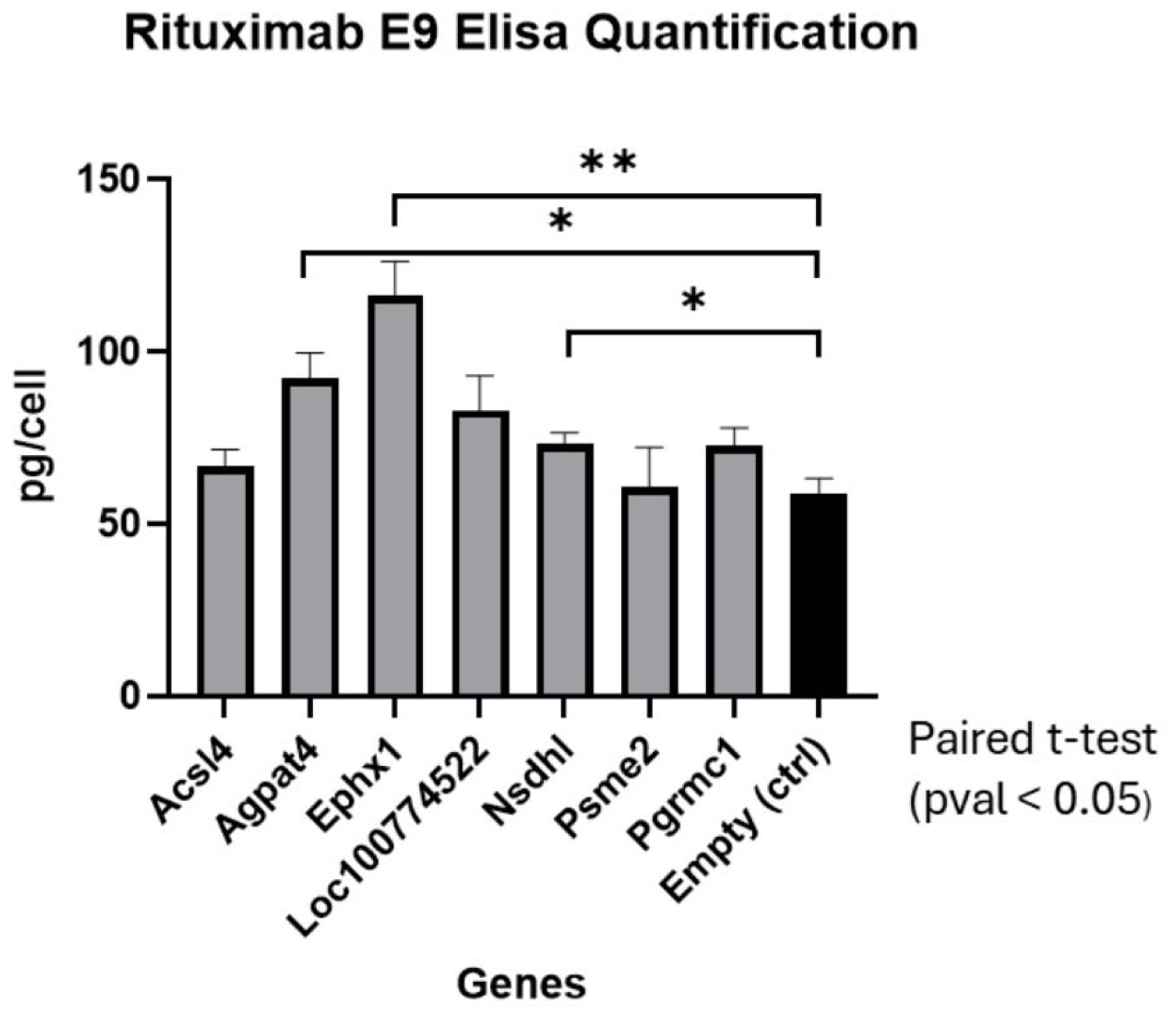
Validation of candidate genes identified from proteomic and transcriptomic datasets using overexpression and Fc-ELISA analysis. **(A)** RNA was extracted two days post-transfection and converted to cDNA for qPCR analysis, confirming successful overexpression of all tested genes in the cells. **(B)** Cell counts and supernatant were collected on day four post-transfection. Indirect Fc-ELISA was performed on the supernatant, revealing that three candidate genes—*Agpat4*, *Ephx1*, and *Nsdhl*—significantly enhanced rituximab secretion in the low-producing cell line, E9, compared to the control (empty vector backbone; paired t-test, p < 0.05).

On day 4, we performed cell counts to determine viability and harvested supernatant for rituximab quantification by ELISA. The obtained concentrations were normalized to viable cell numbers, resulting in their yield (pg/cell) (Figure 5B). Compared to the control (empty vector backbone), overexpression of Agpat4, Ephx1, and Nsdhl significantly improved rituximab secretion in cell line E9 (p < 0.05, paired t-test) with fold changes of 1.57, 1.98, and 1.25 respectively. These three proteins were detected only in rituximab-producing cells in the FcBAR proteomic data and not in the control cell line indicating that these protein interactors are rituximab specific. In addition, paired t-test shows that there was no significant difference (p > 0.05) in cell viability between the overexpression of these validated genes and the control (Supplementary Figure S1B). These proteins all localize to the ER, suggesting that ER-related processes represent a key bottleneck in rituximab secretion in these cells.

## Discussion

Here we presented and used FcBAR to study a secreted protein, rituximab. We target the Fc regions of the rituximab rather than other domains such that the Fc region is highly conserved across antibodies, enabling this method to be extended to other mAbs or Fc-fusion proteins without modification, Fc-directed labeling avoids interference with the antigen-binding site (Fab), reserving antibody function, and Fc-specific detection antibodies are more widely available. We hypothesized that our Fc-targeted approach should capture proximal interactors across the entire antibody structure, as rituximab has a maximum linear dimension of 197 Angstrom (19.7 nm) (Belviso et al., 2022), while the biotinylation radius spans 10 to >200 nm (Rega et al., 2023). Since the labeling radius comfortably exceeds the antibody’s dimensions, we would expect to detect primary Fc interactors (direct binders) and proteins associated with other antibody regions (Fab, hinge) or proximate to the Fc regions. Cells are harvested during the mid-exponential phase for FcBAR as this ensures that the cells are actively growing and maintaining high levels of protein synthesis to capture protein interaction with rituximab before secretion. In contrast, if cells are harvested during the lag phase, the cells are still adapting to their environment and preparing to divide. If harvested during the stationary phase, cell growth slows down, protein products are at its peak of secretion, waste products accumulate, and the cells enter survival mode (González-Hernández & Perré, 2024; Templeton et al., 2013).

Thus, FcBAR allowed us to identify PPIs that enhance secretion through the secretory pathway. We expressed rituximab in suspension CHO-S cells and identified a range of clones with different productivity levels. To identify factors affecting productivity, we employed FcBAR to study the PPIs of rituximab, along with RNA-Seq, integrating both approaches to identify proteins that interact with rituximab in the secretory pathway. We then performed linear regression analysis of the FcBAR data to identify proteins that are associated with high productivity and identified proteins that are differentially expressed in high producers in both the proteomic and transcriptomic datasets. For validation through overexpression, we selected proteins from both analyses that are involved in lipid metabolism, protein transport, and protein modification.

Our validation experiments revealed that overexpression of AGPAT4, NSDHL, and EPHX1 significantly increased rituximab secretion. Notably, two of these proteins are directly involved in lipid metabolism. AGPAT4 (LPATTδ) belongs to the acrylglycerolphosphate acyltransferase (AGPAT) enzyme family, which catalyzes the synthesis of phosphatidic acid (PA) from lysophosphatidic acid (LPA) and acyl-CoA, involved in phospholipid and triacylglycerol biosynthesis (Karagiota et al., 2022). LPA is involved in membrane trafficking, membrane curvature, and protein movement within the cell (Zhukovsky et al., 2019). AGPAT4 is located in the ER, Golgi apparatus, and mitochondria. It interacts with the BAR-domain-containing protein, a member of the C-terminal binding protein (CtBP) family, which plays a role in membrane fission and post-Golgi carrier formation (Pagliuso et al., 2016). Through this interaction, AGPAT4 aids in Golgi membrane fission by converting LPA to PA, which induces membrane curvature or recruits proteins that bind to PA (Karagiota et al., 2022; Pagliuso et al., 2016).

The second lipid-related protein we identified, NSDHL, is a NAD(P)H steroid dehydrogenase-like enzyme involved in cholesterol biosynthesis (Caldas, 2003; Cunningham et al., 2009). High NSDHL expression is associated with highly prolific cells (Cunningham et al., 2009). Located in the ER and associated with lipid droplets, NSDHL plays a role in cholesterol biosynthesis and lipid metabolism. To properly localize to the ER membrane, NSDHL requires transport through the Golgi apparatus (Caldas, 2003).

The third validated protein, EPHX1, indirectly interacts with fatty acids. EPHX1 is a microsomal epoxide hydrolase located in the microsomal fraction of the ER, where it converts highly reactive epoxides into less reactive diols, such as trans-dihydrodiols (Gautheron & Jéru, 2021; Václavíková et al., 2015). EPHX1 plays a role in hydrolyzing endogenous epoxy fatty acids (EpFAs) derived from polyunsaturated fatty acids, such as epoxyeicosatrienoic acids (EETs), which are produced by CYP450 enzymes in the ER (Gautheron & Jéru, 2021). In fibroblasts, EPHX1 knockout prevents adipocyte differentiation, increases oxidative stress, and leads to cellular senescence (Gautheron & Jéru, 2021).

Through this work, we showed that in the study of PPIs and RNA-Seq, we were able to identify new genes that, when overexpressed, increased rituximab production. This may be because both AGPAT4 and NSDHL are lipid-associated proteins whose overexpression might improve lipid and cholesterol metabolism. The expected impacts on membrane composition and fluidity could impact protein trafficking, folding, and secretion, ultimately boosting rituximab secretion. Additionally, overexpression of these genes might activate other pathways that indirectly enhance rituximab production. It would be of interest to determine whether expressing all three proteins or in dual combinations would produce a significant increase in rituximab production compared to individual overexpression, since these proteins have different functions.

Our approach here is a simplified version of BAR, tailored to proteins with an Fc domain. BAR is a versatile protein labeling method that requires only specific binding antibodies for POI, avoiding the laborious process of introducing foreign DNA into the host cell, (Sears et al., 2019). Additionally, BAR can be done on fixed cells and it provides a temporal snapshot of proximal proteins at a specific time point, thus avoiding prolonged labeling times that could affect protein function.

However, BAR does have limitations. The method requires highly specific binding antibodies for the POI, which may not be readily available depending on the organism. Furthermore, BAR does not distinguish between direct and indirect interactions and may miss transient PPIs due to the temporal nature of the labeling process.

Previous applications of BAR have been successfully applied to both non-secreted and secreted proteins. For non-secreted proteins, BAR has been used to study the mitochondrial matrix protein Lamin A/C in HeLa cells, primary cell cultures, and primary human muscle and adipose tissues, successfully identifying putative interactors (Bar et al., 2018). For secreted proteins, BAR has been applied to fetuin-B and able to identify unique interactome corresponding to fetuin-B (Masson et al., 2024a).

In this study, we modified BAR to simplify the protocol for Fc-bearing proteins. We demonstrated it with a model secreted mAb, showing it can identify potential protein interactors of rituximab within the ER and Golgi apparatus. This specificity is possible because HRP is only active in oxidizing environments, such as the lumen of the ER and Golgi apparatus (Bosch et al., 2021).

Taken together, the identification of AGPAT4, EPHX1, and NSDHL as key interacting proteins provides confidence in the utility of FcBAR for gaining valuable insights into optimizing protein secretion in CHO cells. These findings highlight the importance of lipid metabolism and ER function in antibody secretion. Our approach offers a promising strategy for optimizing the manufacturing of complex biologic therapies, potentially reducing production costs and improving patient access. Future studies could explore whether these protein overexpression strategies can be applied to other therapeutic antibodies and difficult-to-express proteins.

## Materials and Methods

### Cell Line Engineering and Protein Expression

The rituximab vector was constructed using the pcDNA3.1(+) backbone with uracil-containing primers and Phusion Hot-Start Flex polymerase (New England Biolabs, Ipswich, MA, USA). The rituximab vector consists of CMV-HC-BGHpA-EF1α-LC-BGHpA with selection marker SV40-NeoR-SV40pA. CHO-S suspension cells (Life Technologies, Carlsbad, CA, USA) were transfected with the rituximab plasmid using FuGENE HD transfection reagent (Promega) as described previously in the supplement (Pristovšek et al., 2019). Cells were selected with 0.500 μg/ml Geneticin Selective Antibiotic (Gibco, USA) to establish a stable polyclonal population. Fluorescence-activated cell sorting (FACS) was used to isolate rituximab-expressing clones, which were subsequently expanded as previously described (Pristovšek et al., 2018). The expanded clones were cultured in 125 mL Erlenmeyer shaker flasks with a baffled bottom (Fisher Scientific) in the presence of 0.25 mg/ml Geneticin.

### Quantifying Rituximab Productivity Through Surface Plasmon Resonance

Rituximab titer was determined for each clone using an Octet RED96 (Pall, Menlo Park, CA, USA, a biolayer interferometry instrument that uses surface plasmon resonance (SPR) to measure protein binding as described previously (Grav et al., 2015), with cell counts taken daily. Following equilibration in sample diluent buffer (PBS with 0.1% BSA and 0.02% Tween-20), Protein A biosensors measured rituximab concentrations for 120 seconds at 30°C. Absolute concentrations were determined by comparing measurements to a calibration curve prepared from human IgG dilution series (GENSA01006, VWR, Dublin, Ireland). Biosensor tip regeneration was performed between measurements using 10 mM glycine (pH 1.7). Specific productivity (pg/cell/day) for rituximab clones was calculated based on rituximab titer and the integral viable cell density measurements as previously described (Pybus et al., 2013) to determine high and low rituximab producing clones.

### Cell Culture

A panel of ten cell CHO-S clones expressing rituximab were obtained. A Laminin-alpha1 knock-out CHO-S clone line was obtained to use as a control. CHO-S cells were grown in 125ml Erlenmeyer shaker flasks with baffled bottom (Fisher Scientific, USA) in a humidified 37°C incubator with shaking at 130 RPM at 5% CO2 with CD CHO Medium (Gibco, USA) supplemented with 8mM L-Glutamine (Gibco) and 1x anti-clumping agent (Gibco). The rituximab producing clones were grown in media containing 250 ug/ml Geneticin Selective Antibiotic (G418 sulfate; Gibco) for selection.

### Western blot

Western blots were performed to analyze rituximab using a precast polyacrylamide gel, 4-15% Mini-Protean TGX Gels (Bio-Rad, Hercules, CA, USA). The samples were run under reducing conditions by mixing with loading dye (4x Laemli + 10% β-Mercaptoethanol) (Bio-Rad) and boiling at 95°C for 5 minutes. The gel was run on a Mini-PROTEAN Tetra Vertical Electrophoresis Cell (Bio-Rad) at 160V for 40 minutes with the Precision Plus Protein standards ladder (Bio-Rad). The gel was transblotted onto 0.2 μm nitrocellulose membranes (Bio-Rad) with the Trans-Blot Turbo Transfer System (Bio-Rad). The membranes were blocked with Intercept Blocking Buffer (LI-COR, Lincoln, NE, USA) for 1 hr. Rituximab was detected with IRDye 800CW Goat anti-human IgG Secondary Antibody (LI-COR) at 1:15,000 in Intercept Antibody Diluent (LI-COR) for 1.5 hours at room temperature. The membranes were washed 3x for 10 minutes with 0.1% TBST and imaged using the LI-COR 9120 Odyssey Infrared Imaging System.

For staining of intracellular biotinylated proteins, 20 μg of total protein from labeled cells was loaded and transblotted. The membrane was blocked by 3% BSA in 0.1% TBST for 1 hour and probed with HRP-conjugated streptavidin (Abcam, Cambridge, UK) diluted in blocking buffer at 1:2000 for 40 minutes. For visualizing the proteins’ bands, the Clarity Western ECL Substrate (Bio-Rad) was used.

### Quantifying HC and LC with qPCR

Total RNA was extracted from 1-2 × 10^6^ cells using the RNeasy Plus Mini Kit (Qiagen, Hilden, Germany) according to the manufacturer’s instructions. RNA concentration was measured with Qubit fluorometric analysis (Life Technologies,Carlsbad, California, USA). cDNA was synthesized from 1.5-3 µg of total RNA using the Maxima First Strand cDNA Synthesis Kit for RT-qPCR with dsDNAse treatment (Thermo

Fisher Scientific). RT-qPCR analyses were performed on the QuantStudio 5 Real-Time PCR System using TaqMan^TM^ Multiplex Master Mix (Thermo Fisher Scientific) in triplexes (gene of interest and two normalization genes) using the following amplification conditions: 50°C for 2 min, 95°C for 10 min; 40x: 95°C for 15 s, 60°C for 1 min. Custom-made TaqMan assays were used for rituximab light chain and heavy chain, as well as normalization genes Gnb1 and Fkbp1a (Brown et al., 2018). All the primers and probes were validated by melting curve analysis and primer efficiency test. Using the ΔΔCt method, the relative expression levels of rituximab heavy and light chains were calculated by normalization to the geometric mean of expression levels of the two normalization genes. Each experiment included controls with no template and was performed using technical triplicates.

### Quantification of PPIs of Rituximab in CHO-S cells using the FcBAR method

FcBAR was used to target rituximab in 10 CHO-S clones that express rituximab at different productivity levels, in order to identify PPIs in each clone. To accomplish this, CHO-S clones expressing rituximab and non-producing CHO-S control cells, each in technical triplicate, were harvested at mid-exponential phase, fixed in 4% paraformaldehyde (PFA) in PBS (Thermo Scientific), and permeabilized with 0.4% PBST. Endogenous peroxidases were inactivated with 0.4% H[O[(Fisher Scientific, USA), and cells were blocked with 5% goat serum (Gibco) in 1% BSA-PBST. Cells were then incubated with goat anti-human IgG Fc secondary antibody conjugated to HRP (Invitrogen, USA) for 1 hour at room temperature with rotation. Proximal protein biotinylation was performed by treating cells with H[O[and tyramide-biotin using the TSA Biotin Reagent Pack (SAT700001EA, Akoya Biosciences, Marlborough, MA, USA), resulting in tyramide-biotin radicalization and deposition onto proximal proteins. The reaction was quenched with 0.5 M sodium ascorbate in PBS (Sigma-Aldrich). Cell lysates were extracted using 1.5% SDS (G Biosciences, St. Louis, MO) and 1% sodium deoxycholate (bioWorld, Dublin, OH, USA) in 0.1% PBST, then heated at 99°C for 1 hour with mild shaking. Samples were centrifuged at maximum speed for 5 minutes, and the supernatant was collected for protein quantification using the BCA Protein Assay (Lambda Biotech, Inc., St. Louis, MO, USA). Samples were stored at −80°C until shipment to the Sanford Burnham Prebys Proteomics Core (San Diego, CA, USA) for LC-MS/MS analysis. Biotinylated proteins were enriched using streptavidin beads, digested with trypsin, and subjected to LC-MS/MS analysis.

### Fluorescent Microscopy

Co-localization of rituximab and biotinylated proteins in the endomembrane system was assessed by fluorescent staining and confocal super-resolution microscopy following the BAR protocol. For co-localization studies, a subsample of labeled cells was probed with goat anti-human-DyLight 650 conjugate (Thermo Fisher Scientific) and streptavidin-DyLight 594 conjugate (Thermo Fisher Scientific) to target rituximab and biotinylated proteins, respectively. These antibodies were diluted 1:300 and 1:1000 in the blocking buffer, respectively, and incubated with fixed cells for 30 minutes at room temperature. Cells were washed, counterstained with DAPI, and mounted on slides using antifade Vectashield mounting medium (VectorLabs, Newark, CA, USA). Images were acquired using a Leica SP8 confocal microscope with Lightning deconvolution (Wetzlar, Germany). Colocalization analysis was performed on deconvolved images using the Coloc_2 plugin in Fiji (ImageJ 1.52p) to assess pixel intensity overlap between the 650 nm channel (rituximab) and 594 nm channel (streptavidin) (Schindelin et al., 2015). This analysis generates Pearson’s correlation coefficients (range: −1.0 to 1.0) to quantify the degree of colocalization between the two fluorescent signals. Background subtraction was applied using Fiji’s rolling ball algorithm within defined regions of interest, and threshold values were automatically determined using Coloc_2’s bisection method to optimize background correction.

### LC/MS-MS on Biotinylated Samples

The BAR samples were sent to Sanford Burnham Prebys Proteomic Core for LC-MS/MS analysis. Biotinylated proteins were affinity-purified using the Bravo AssayMap platform (Agilent) with AssayMap streptavidin cartridges (Agilent). On-cartridge digestion of the bound proteins was performed using mass spectrometry-grade Trypsin/Lys-C Rapid digestion enzyme (Promega, Madison, WI) at 70°C for 2 hours. The resulting peptides were desalted on the Bravo platform using AssayMap C18 cartridges. Organic solvents were removed using a SpeedVac concentrator prior to LC-MS/MS analysis. The dried peptides were reconstituted in 2% acetonitrile with 0.1% formic acid and analyzed via LC-MS/MS using a Proxeon EASY nanoLC system (Thermo Fisher Scientific) coupled with a Q-Exactive Plus mass spectrometer (Thermo Fisher Scientific).

Peptide separation was conducted on an analytical C18 Aurora column (75µm x 250 mm, 1.6µm particles; IonOpticks) at a flow rate of 300 nL/min. The 80-minute gradient applied was: 0% to 6% buffer B over 0.5 minutes, 6% to 23% B over 50 minutes, 23% to 34% B over 29 minutes, and 34% to 48% B over 0.5 minutes (Buffer A: 0.1% FA; Buffer B: 80% ACN with 0.1% FA). The mass spectrometer operated in positive data-dependent acquisition mode. MS1 spectra were acquired at a resolution of 70,000, with an AGC target of 1e6 and a mass range of 350-1700 m/z. Up to 12 MS2 spectra per duty cycle were triggered, fragmented using HCD, and acquired at a resolution of 17,500, with an AGC target of 5e4, an isolation window of 1.6 m/z, and a normalized collision energy of 25. Dynamic exclusion was enabled with a duration of 20 seconds.

Mass spectra were analyzed using MaxQuant software (Tyanova, Temu, & Cox, 2016), version 1.5.5.1. MS/MS spectra were searched against the *Cricetulus griseus* Uniport (version August 2020) and GPM cRAP sequences (common protein contaminants). Precursor mass tolerance was set to 20 ppm for the initial search, which included mass recalibration, and 4.5 ppm for the main search. Product ions were searched with a mass tolerance of 0.5 Da. The maximum precursor ion charge state for the search was set to 7. Carbamidomethylation of cysteines was included as a fixed modification, while oxidation of methionines and N-terminal acetylation were considered variable modifications. The enzyme specificity was set to trypsin, allowing up to two missed cleavages. The target-decoy-based false discovery rate (FDR) filter for spectrum and protein identification was set to 1%.

### FcBAR Analysis

The FcBAR proteomic mass spectrometry dataset was preprocessed and analyzed using the DEP package (Zhang et al., 2018) in RStudio to identify differentially expressed proteins. Proteins were assigned unique names, and samples were filtered to remove proteins with excessive missing values. Specifically, proteins were retained only if they were identified in at least 2 out of 3 replicates for any given condition. Samples were normalized using variance stabilizing transformation (VSN), and missing values were imputed based on data missing not at random (MNAR) using a manually defined left-shifted Gaussian distribution. The data were then visualized using Principal Component Analysis (PCA) of the normalized and imputed data. Proteins were filtered using the following criteria: detected in ⅔ replicates for each producer clone, present in <2 replicates in control, or present in ⅔ replicates in control but having a fold change >1 and adjusted p-value (p-adj) < 0.05. These were marked as potential target proteins. Welch’s t-test was performed comparing high producer cell lines versus low producer cell lines to identify proteins that were significantly expressed with p < 0.05 and fold change of >0 in the high producer cell lines. Linear regression analysis was performed in RStudio on the FcBAR proteomic data, examining the relationship between the mean expression of each protein and the productivity level (pg/cell/day) from each rituximab clone.

### RNA-Seq Prep and Analysis

The rituximab clones and its parental cell line were each cultured in triplicate. They were each harvested at the same time as the sample used for FcBAR. Total RNA was extracted from each sample using the RNeasy Kit (Qiagen), following the manufacturer’s instructions, and then quantified using nanodrop. Samples were sent to UCSD IGM Genomics Center for mRNA library preparation and sequencing. RNA quality was analyzed using the Agilent Tapestation 4200, and only samples with an RNA Integrity Number (RIN) exceeding 8.0 were selected for library construction. Libraries were prepared using the Illumina Stranded mRNA Sample Prep Kit with Illumina RNA UD Indexes (Illumina, San Diego, CA), following the manufacturer’s guidelines. The finalized libraries were multiplexed and sequenced on an Illumina NovaSeq X Plus platform, generating 150 base pair (bp) paired-end reads (PE150) with an approximate depth of 25 million reads per sample. The samples were quality controlled using FastQC and the adapter sequences trimmed using Trimmomatic. The sequences were demultiplexed with bcl2fastq v2.20 Conversion Software. Reads were mapped to the CHO GCF_003668045.1_CriGri-PICR genome, aligned and quantified with Salmon to obtain TPM values. Statistical analysis was done in RStudio.

### Generating Plasmids for Overexpression

The Acsl4, Agpat4, Ephx1, Loc100774522, Nsdhl, Psme2, Pgrmc1, and the empty vector backbone (pcDNA3.1(+)) plasmids were designed from Chinese amster (*Cricetulus griseus*) and purchased from GenScript (Piscataway, NJ, USA). The plasmids were transformed into DH5[Competent Cells (Invitrogen) and expanded in Miller’s LB Broth (Corning Inc., Corning, NY, USA) supplemented with 100 μg/ml of ampicillin (Sigma-Aldrich). The plasmids were extracted using QIAprep Spin Miniprep Kit (Qiagen).

### Transient Transfection

PEI Max transient transfection was used for overexpression of the validation genes. PEI MAX-Transfection Grade Linear Polyethylene Hydrochloride (MW 40,000) (Kyfora Bio, Warrington, PA, USA) was dissolved at 1 mg/ml in distilled water and pH was adjusted to 7.0 using 1M of NaOH. The dissolved PEI was sterile filtered in 0.22 μm steritop (MilliporeSigma, Burlington, MA, USA). A day before transfection, cells were passaged at 0.5-0.8 × 10^6^ cells/ml in media containing no anti-clumping. On the day of transfection, cell counts were done and were at least > 95% viable for transfection. Cells were harvested and resuspended at 1.0 × 10^6^ cells/ml in media containing no anti-clumping. Cells were transfected with 1 μg plasmid and 7 μg of PEI Max solution per ml of cell culture (5% of total culture volume). The DNA/PEI mixture was incubated in OptiPRO SFM (Life Technologies) for 20 minutes at room temperature before adding to the cells in 6-well non-treated, polystyrene, flat bottom wells plate (Genesee Scientific). The cells were shaken at 130 RPM at 37°C in 5% of CO_2_. One day post-transfection, 1× of anticlumping agent (Gibco) and 0.6 mM valproic acid sodium salt (Sigma-Aldrich) were added to each well. Some cells were harvested on day 2 post transfection for RNA extraction. Viability was performed on day 4 post-transfection and supernatant was harvested.

### qPCR to Confirm Gene Overexpression

RNA extraction was done day 2 post transfection following the RNeasy Mini kit protocol (Qiagen) to confirm overexpression of the validation genes. RNA was converted to complementary DNA (cDNA) following the SuperScript II RT protocol (Thermofisher) with 1 μg of RNA and RNAseOUT Recombinant Ribonuclease Inhibitor (Life Technologies). Quantitative PCR (qPCR) was performed using iTaq™ Universal SYBR Green Supermix (Bio-Rad) using primers targeting the validating genes purchased from IDT DNA (USA) (Table 1) with Gnb1 as the housekeeping gene in 96-well PCR Plates (Bio-Rad). The samples were run as follows: 95°C for 2 min; 40×: 95°C for 10 sec, 60°C for 30 sec; 65°C for 5 sec, 95°C for 5 sec. Using the ΔΔCt method, the relative expression levels of each gene were calculated by normalization to the expression level of the normalization gene. Each experiment included no-template controls and was performed using technical duplicates.

**Table 1:**
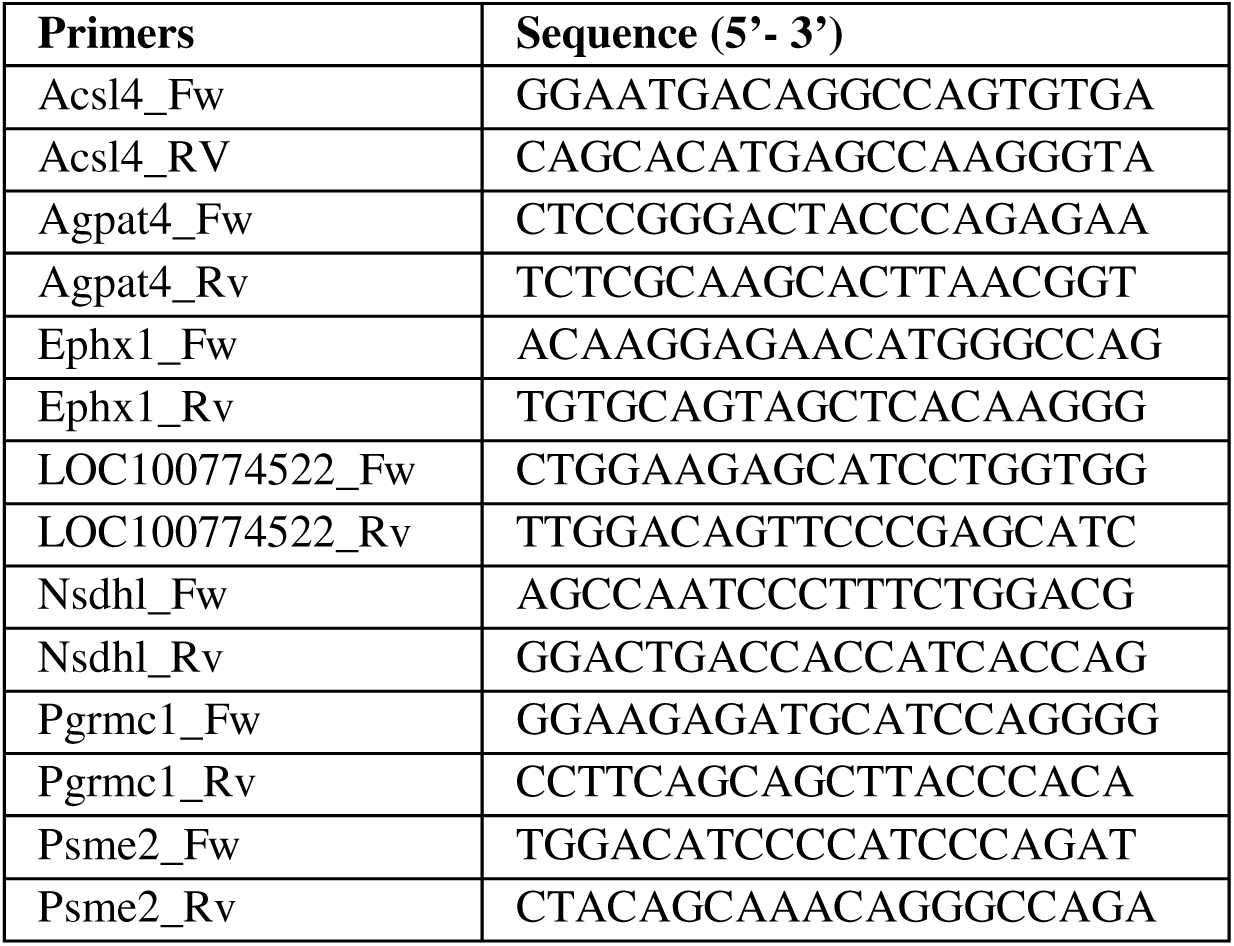
Primers of validating gene.

### Fc-ELISA

Rituximab secretions (supernatant) were quantified by ELISA on day 4 post transfection of the validation genes. Ninety-six-well microplates (Maxisorp; Thermo Fisher) were coated with 2 μg/ml of Goat Anti-Human IgG (I1866; Sigma-Aldrich) in coating buffer (Elisa Reaction Buffer, Carbonate-Bicarbonate, 50 mM; BioWorld) and incubated at 4°C overnight. After the plates were washed 3× with 0.05% PBST, the plates were blocked with 1% BSA in PBS for 2 hours at 37°C. The plates were washed 3× with 0.05% PBST. The supernatant was tested in duplicate at 1:600 along with the standard IgG human serum (I4506; Sigma-Aldrich) in blocking buffer. The samples were incubated at 37°C for 2 hrs. After washing 3× with 0.05% PBST, the samples were incubated with anti-human IgG (Fc specific) HRP (A0170; Sigma-Aldrich) at 1:5,000 in PBS for 1 hour at 37°C. The plates were washed 3× with 0.05% PBST before adding TMB substrate (T8665; Sigma-Aldrich) and incubated for 4-5 minutes. The reaction was stopped with 1M sulfuric acid. The absorbance was read at 450 nm and 595 nm using the BioTek Synergy MX microplate reader (Agilent, Santa Clara, CA, USA). The concentrations were calculated by curve fitting in GraphPad Prism software with hyperbola.

### Statistical Analyses

Statistical analysis of BAR proteomic data was performed using the DEP package (version 1.24.0) in RStudio (version 4.3.3) and Microsoft Excel. RNA sequencing data was analyzed using the DESeq2 package (version 1.42.1) and clusterProfiler package (version 4.10.1) in RStudio, with genome annotation provided by the org.Hs.eg.db package (version 3.18.0). Data visualization included volcano plots, RNA-Seq PCA plots, and enrichment pathway plots, which were generated using the ggplot2 package (version 3.5.0). BAR proteomic PCA plots were created with the DEP package. Bar graphs for qPCR and validated gene expression data were generated using GraphPad Prism using paired t-test (version 10.4.0). All figures were designed and assembled using BioRender.com.

## Acknowledgements

This work was supported by generous funding from NIH (R35 GM119850), NSF (CBET-2030039), and the Novo Nordisk Foundation (NNF20SA0066621). The authors would also like to thank Henning Gram Hansen for his support in generating the rituximab clones.

## Data Availability

Study findings are supported by data available upon request from the corresponding author. Mass spectrometry proteomics data, raw and processed, has been uploaded to theMassIVE (member of the proteome Xchange consortium) repository at https://doi.org/10.25345/C5ZZ0V, reference number PXD063116. Raw and processed transcriptomic data has been uploaded to GEO with the data set identifier [GSE295486].

## Supplementary Figures and Tables

**Supplementary Figure S1:**
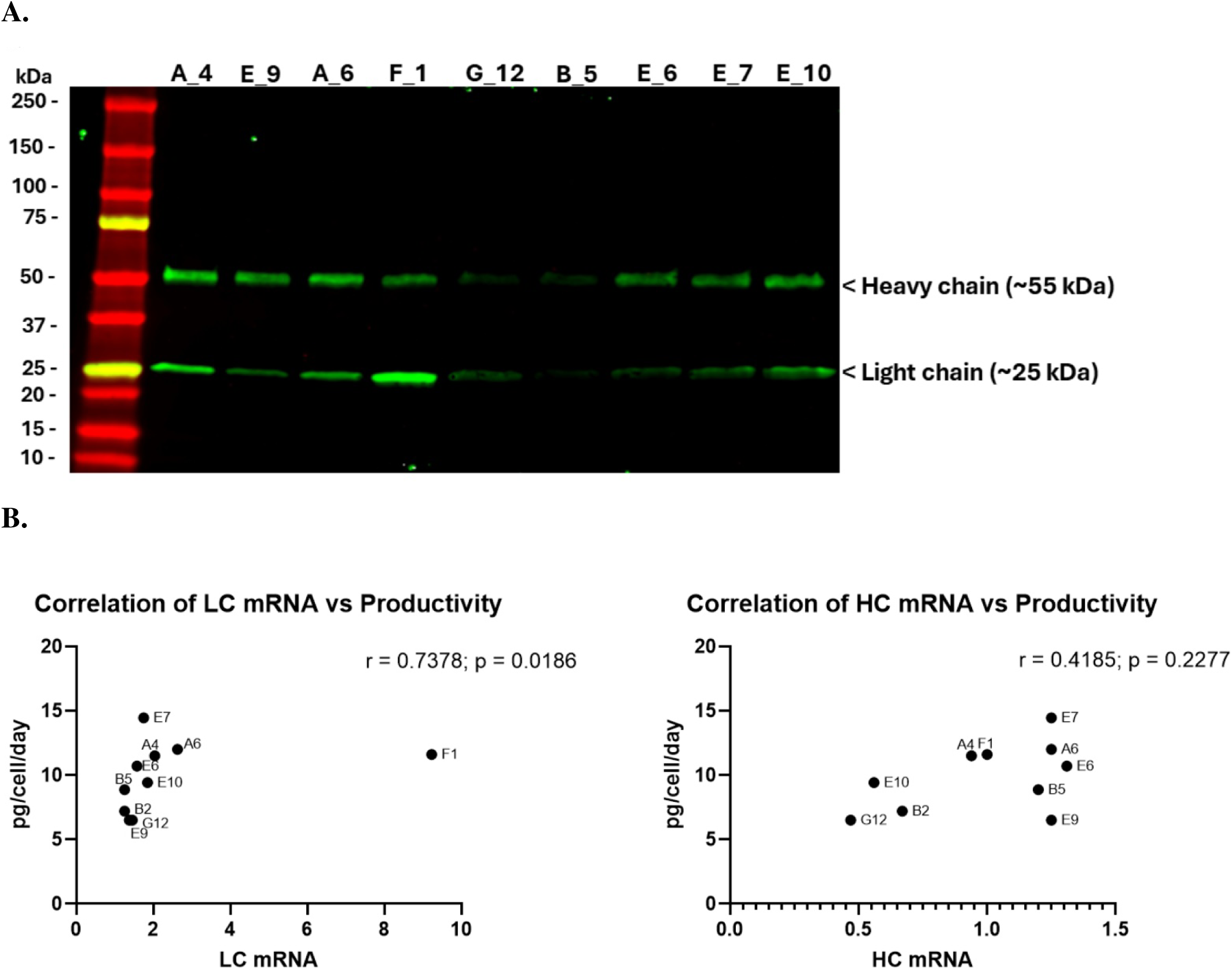
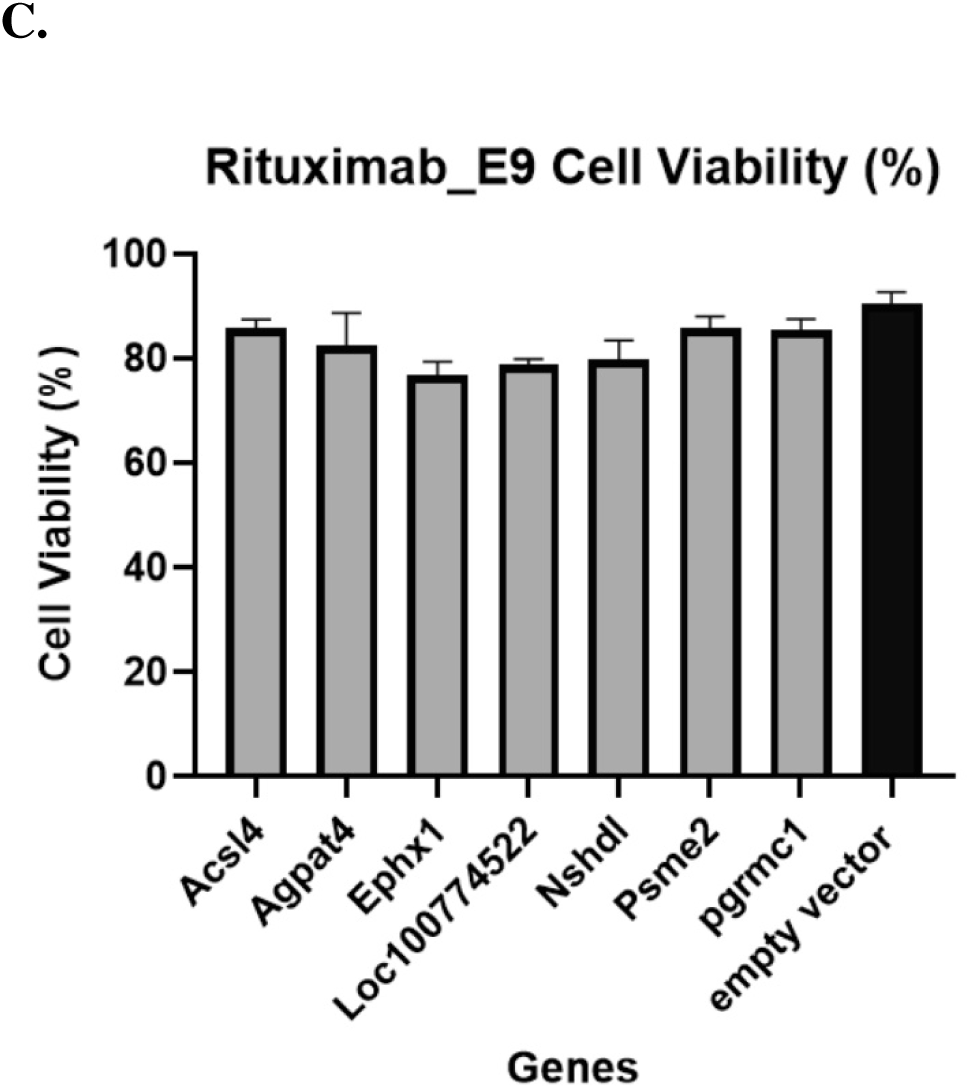
Rituximab clone secretion confirmation by Western blot, correlation of heavy chain and light chain rituximab expression with productivity, and rituximab E9 overexpression viability post-transfection. **(A)** Rituximab secretion from clone supernatants was confirmed using Western blot analysis. The membrane was stained with IRDye 800CW Goat-anti-Human IgG Secondary Antibody and imaged with a LI-COR infrared (IR) scanner, confirming rituximab secretion and demonstrating the expected light chain (∼25 kDa) and heavy chain (∼55 kDa) bands. Rituximab clone B2 was not included. **(B)** qPCR was performed targeting the heavy chain (HC) and light chain (LC) of rituximab. Productivity (pg/cell/day) was determined with SPR. Spearman rank correlation analysis between HC expression and productivity revealed a low positive correlation that was not statistically significant (r = 0.4185, p = 0.2277). In contrast, a significant correlation was observed between LC expression and productivity, with an outlier identified among the cell lines (r = 0.7378, p = 0.0186). **(C)** Cell count and viability were assessed on day 4 post-transfection with the validation genes. Paired t-test analysis showed no significant difference (p > 0.05) in cell viability between overexpression of the validated genes and the control (empty vector backbone).

**Supplementary Table S1:**
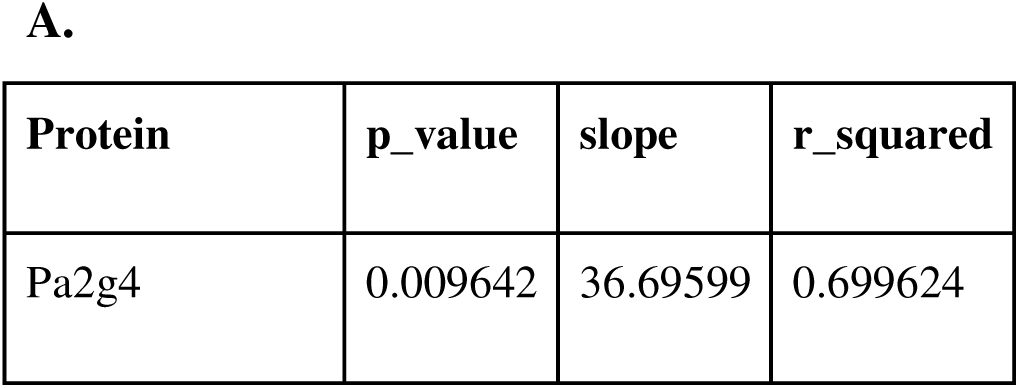
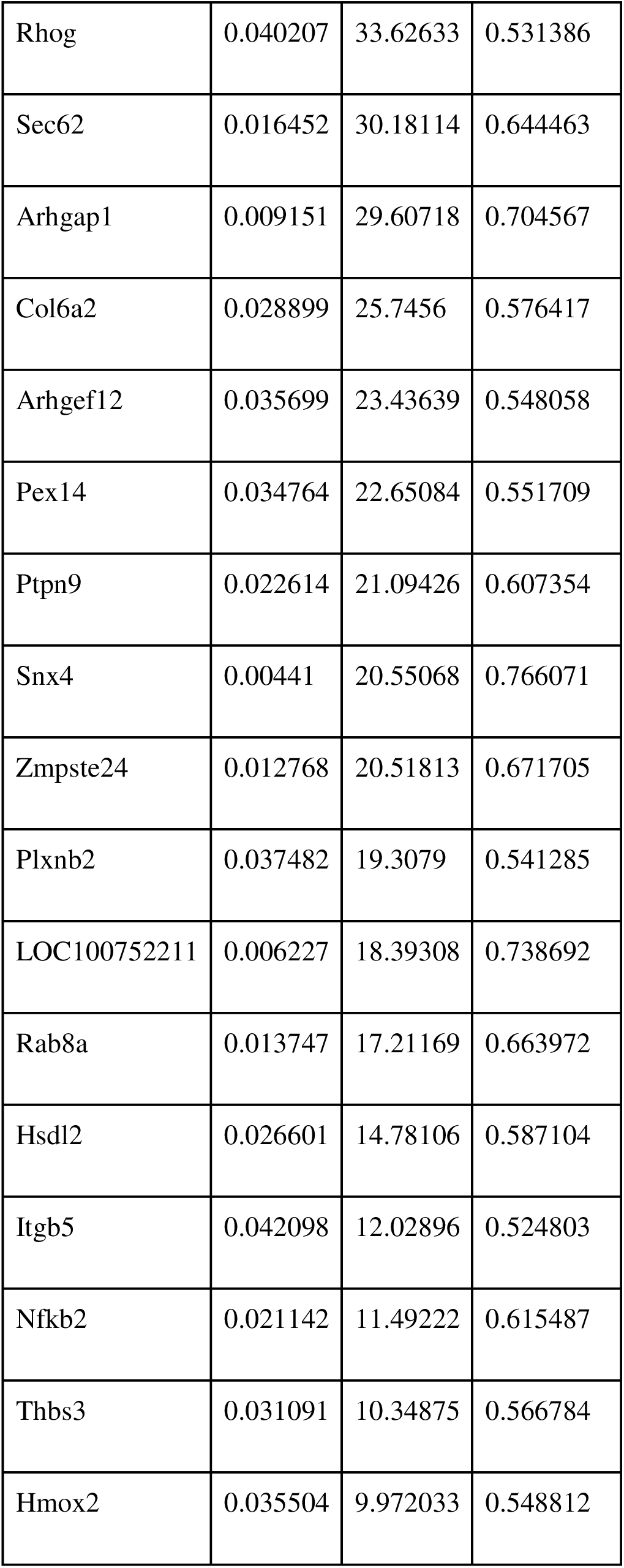
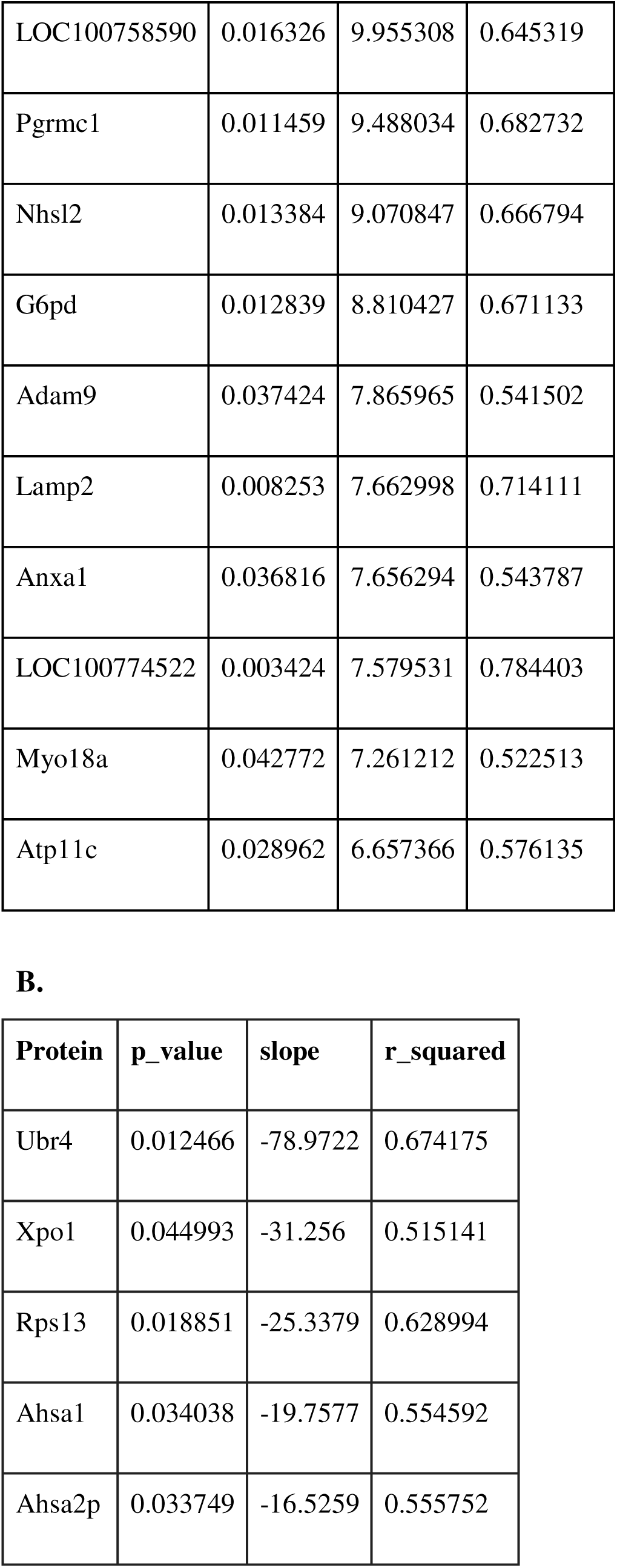
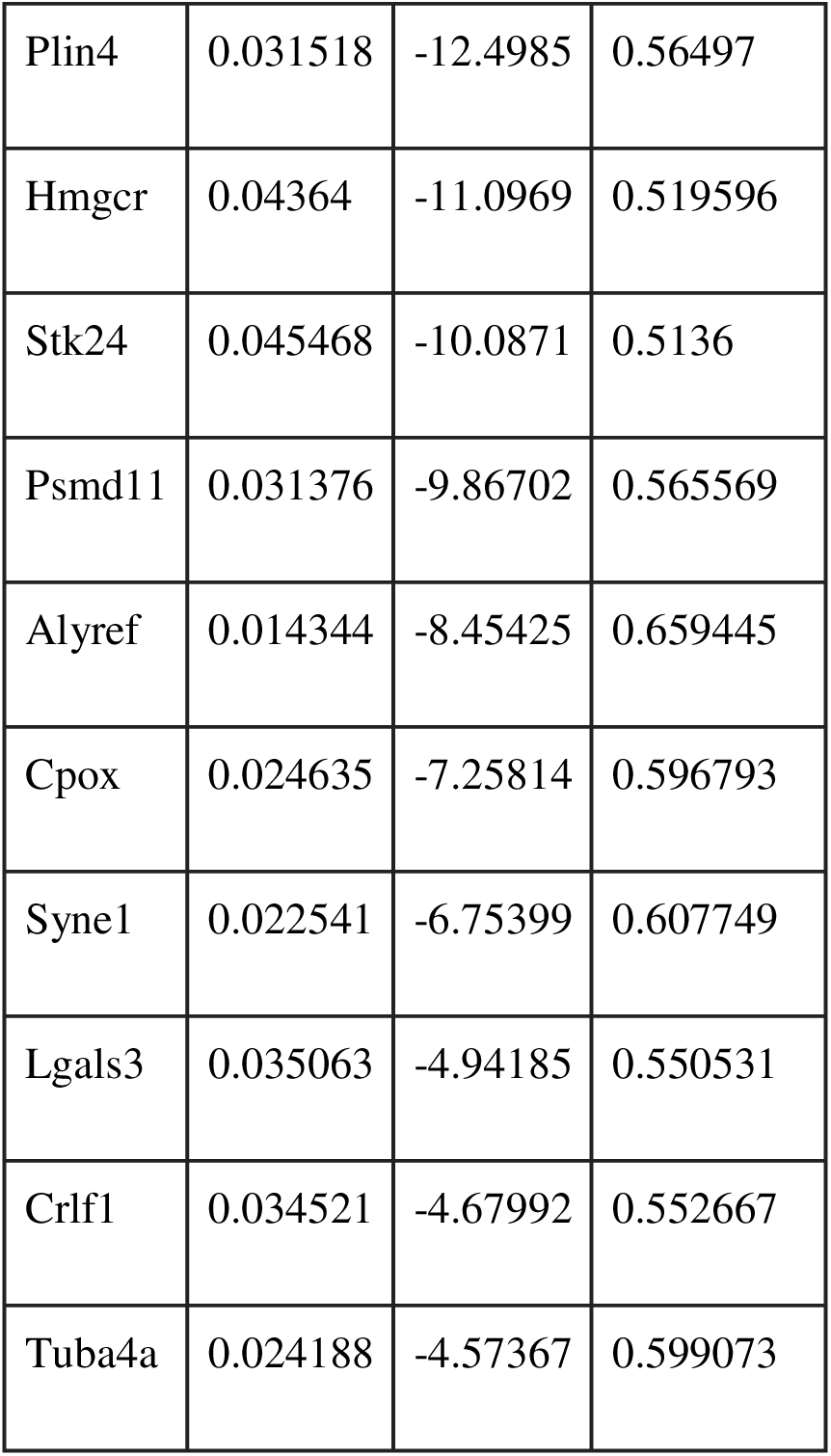
Significant proteins identified through linear regression analysis of mean protein expression versus productivity. Significant proteins that **(A)** positively and **(B)** negatively correlated with productivity (pg/cell/day) in the BAR proteomic dataset. Unfiltered linear regression analysis results for rituximab clones versus productivity are provided in Supplementary Table S3.

**Supplementary Table S2:**
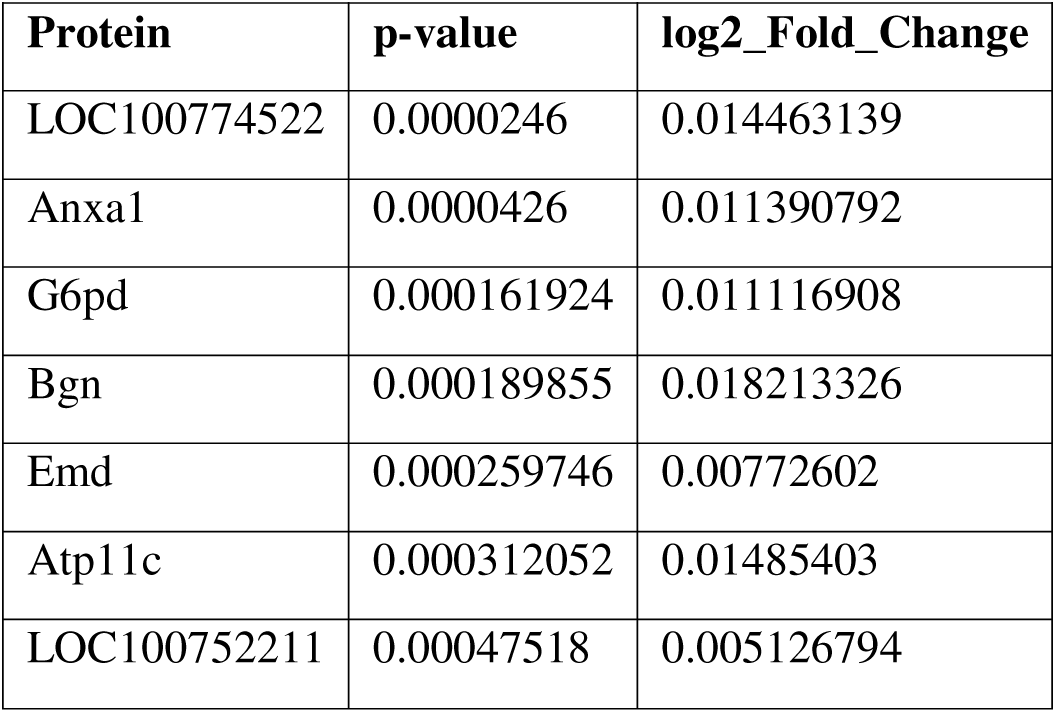
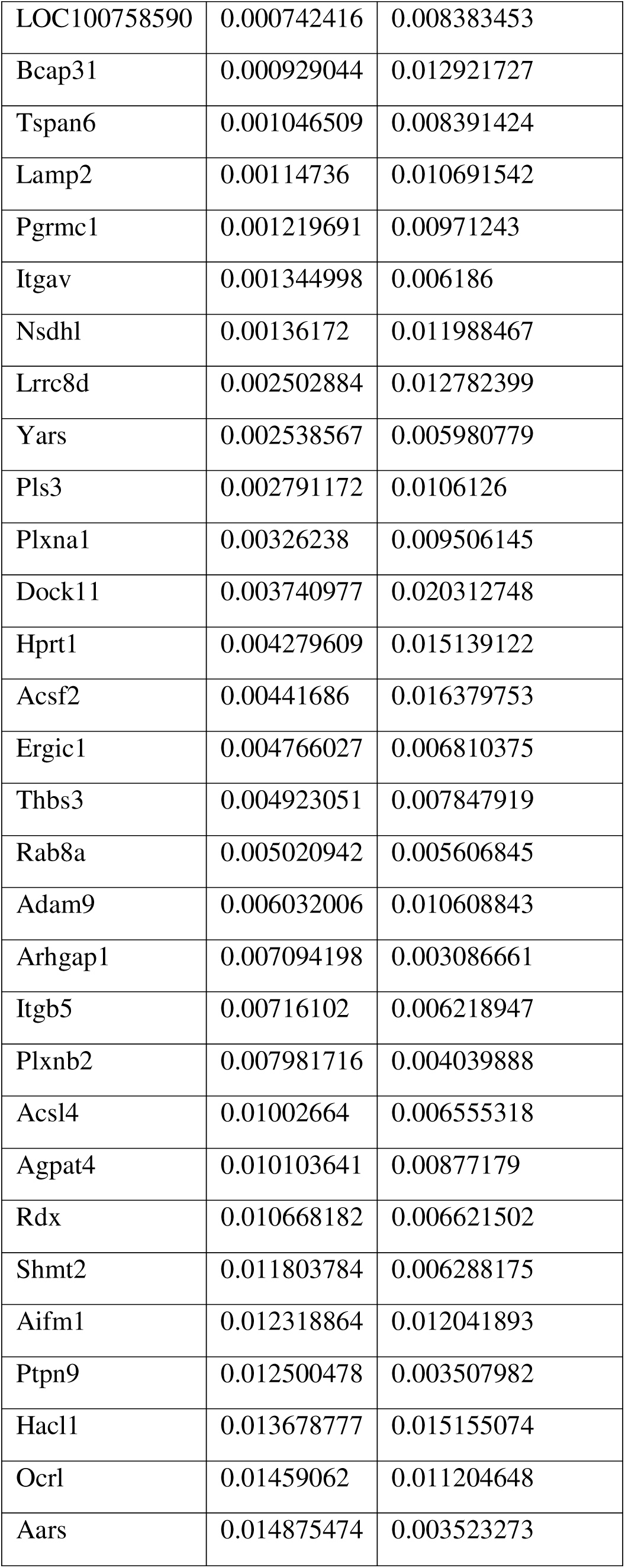
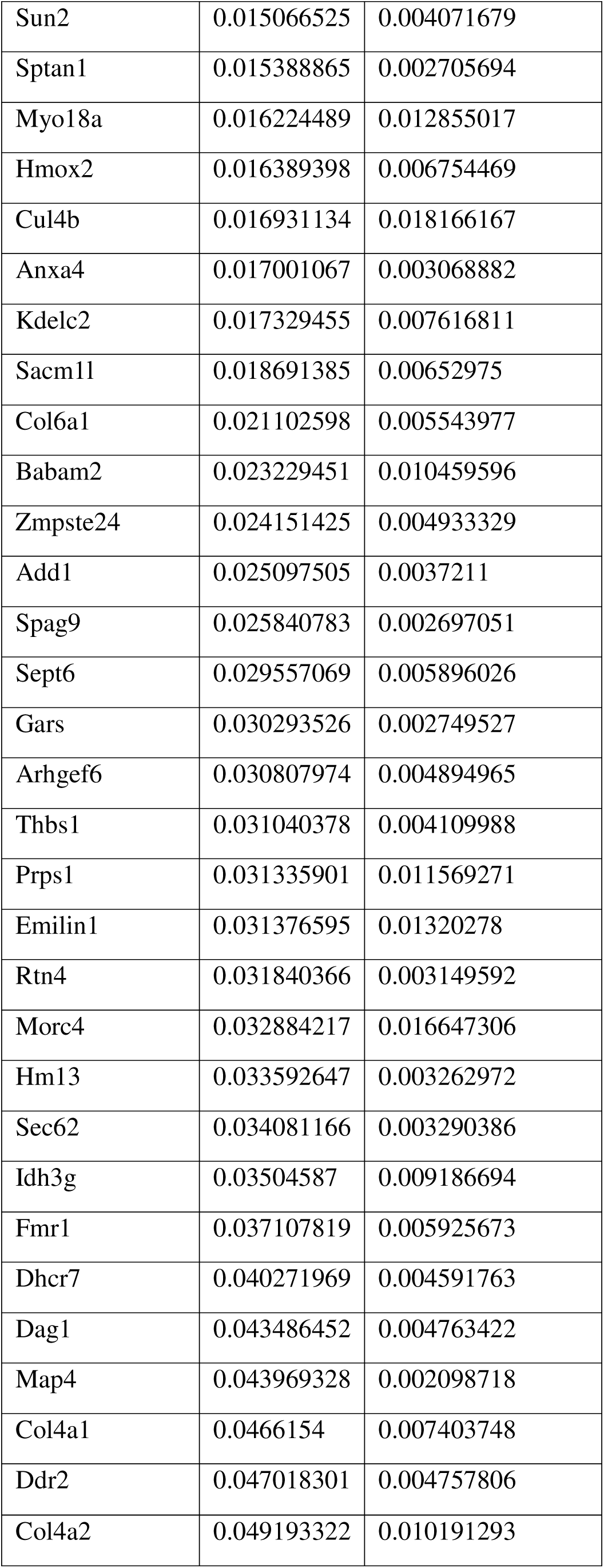
Significant proteins identified in high-producing clones that were consistently detected in both the BAR proteomic and transcriptomic datasets. The table includes associated *p*-values and log□ fold change values corresponding to the proteomic data. Unfiltered p-values and log2 fold changes for high and low rituximab producers in the FcBAR and transcriptomic data are in Supplementary Tables 4 and 5, respectively.

